# Biomolecular Simulations under Realistic Macroscopic Salt Conditions

**DOI:** 10.1101/226001

**Authors:** Gregory A. Ross, Ariën S. Rustenburg, Patrick B. Grinaway, Josh Fass, John D. Chodera

## Abstract

Biomolecular simulations are typically performed in an aqueous environment where the number of ions remains fixed for the duration of the simulation, generally with either a minimally neutralizing ion environment or a number of salt pairs intended to match the macroscopic salt concentration. In contrast, real biomolecules experience local ion environments where the salt concentration is dynamic and may differ from bulk. The degree of salt concentration variability and average deviation from the macroscopic concentration remains, as yet, unknown. Here, we describe the theory and implementation of a Monte Carlo *osmostat* that can be added to explicit solvent molecular dynamics or Monte Carlo simulations to sample from a semigrand canonical ensemble in which the number of salt pairs fluctuates dynamically during the simulation. The osmostat reproduce the correct equilibrium statistics for a simulation volume that can exchange ions with a large reservoir at a defined macroscopic salt concentration. To achieve useful Monte Carlo acceptance rates, the method makes use of nonequilibrium candidate Monte Carlo (NCMC) moves in which monovalent ions and water molecules are alchemically transmuted using short nonequilibrium trajectories, with a modified Metropolis-Hastings criterion ensuring correct equilibrium statistics for an (Δ*µ, N, p, T*) ensemble. We demonstrate how typical protein (DHFR and the tyrosine kinase Src) and nucleic acid (Drew-Dickerson B-DNA dodecamer) systems exhibit salt concentration distributions that significantly differ from fixed-salt bulk simulations and display fluctuations that are on the same order of magnitude as the average.

## Introduction

Molecular dynamics simulations have proven themselves a powerful tool for studying the structure, dynamics, and function of biomolecular systems in atomic detail. Current state-of-the-art approaches simulate a small volume around the biomolecule using explicit atomistic solvent to model the local environment. To more realistically emulate electrostatic screening effects in the local solvent environment, explicit ions are generally added, both to achieve net neutrality and to mimic the macroscopic salt concentration in the *in vitro* or *in vivo* environment being studied.

Salt concentrations and ionic composition are tightly regulated in biology^4^. Ion composition differs between inter/intracellular environments^2^, tumor microenvironments^5^, and organelles^1^ (see Figure 1, *left*). The local ionic concentration in the environment around real biological macromolecules, however, can significantly deviate from macroscopic concentrations. Many biomolecules possess a significant net charge, and the energetic penalty for physical systems to maintain charge separation over large distances serves to recruit more or less ions from bulk to maintain charge neutrality over macroscopic lengthscales. Yet, the number of ions within the immediate vicinity may not necessarily counter the net charge of the macromolecule, as proteins can predominantly bind to ions that have the same polarity as their net charge^6^. Additionally, statistical fluctuations in the total number of ions in the region around the biomolecule may result in significant variance in the local salt concentration, where relative concentration fluctuations diminish slowly with increasing simulation volume (Figure 1, *middle*).

**Figure 1.**
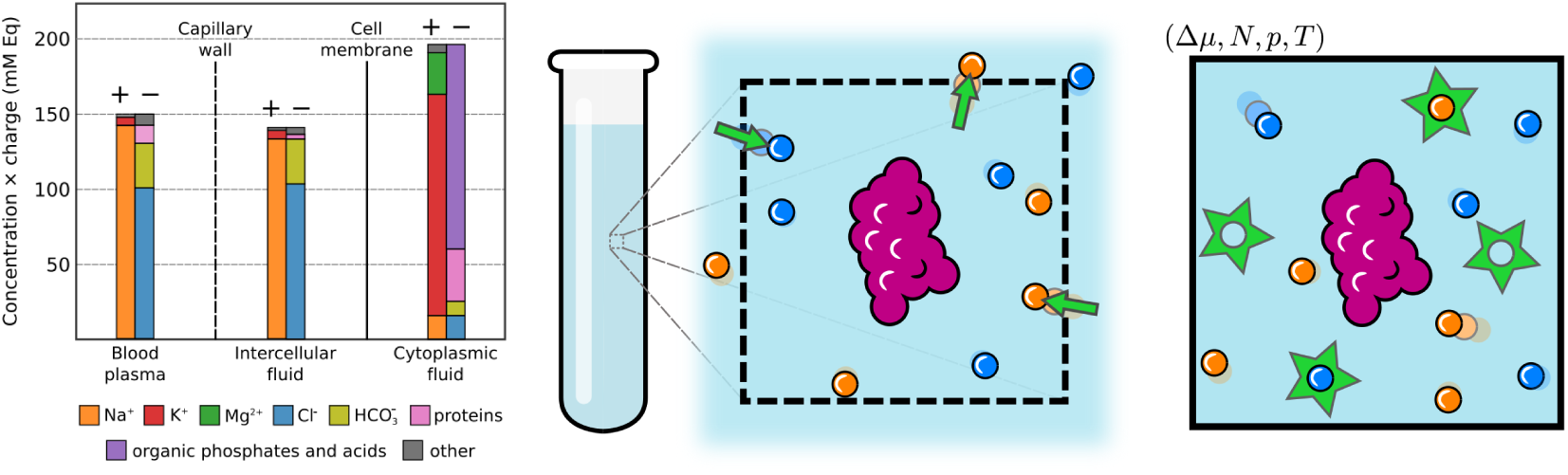
Schematic illustrations of typical salt concentrations in mammalian environments and anions and cations being exchanged with a saline buffer in the region around a biomolecule. *Left:* The ion compositions of intra- and intercellular mammalian environments are shown as *millimolar equivalents* (mM Eq), which is the ion concentration multiplied by the absolute charge of the ion. The primary contribution to the ionic-strength are monovalent ions (Na^+^, K^+^, Cl^−^), divalent cations (predominantly Mg^2+^), complex salt and buffer molecules, and charged proteins. In addition to the significant difference between the ionic composition of the cytoplasmic fluid and extracellular fluid, organelles can also have markedly different ionic concentrations to the cytoplasmic fluid^1^. Over large lengthscales, environments are approximately electrostatically neutral; electrostatic potentials across cell membranes are maintained by an imbalance of anions and cations that is minuscule relative to the total number ions^2^. Figure adapted from^2^ and^3^. *Middle:* In a very large system, where the number of water molecules and number of ions are fixed, significant fluctuations can occur in the ionic strength of the local environment of a biomolecule (in purple). The local environment is represented by a dashed line, within which the number of water molecules and ions fluctuate at equilibrium. *Right:* A simulation with an osmostat replicates the natural variations in ionic strength around a biomolecule that would occur if the system were embedded in an infinite saline reservoir at a fixed macroscopic salt concentration. Anions and cations (blue and orange spheres) are inserted and deleted (green stars) from the system using semigrand canonical Monte Carlo moves that exchange explicit water molecules for the ions in a manner that maintains total charge neutrality. The reservoir is completely defined by its thermodynamic parameters, which in this case include the difference in the chemical potential for two water molecules and NaCl, 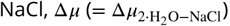, pressure *p*, and temperature, *T*.

### Biomolecular behavior can be sensitive to salt environments

The conformations, dynamics, function, and binding of biological macromolecules can be exquisitely sensitive to the salt concentration and composition of the local environment. The Hofmeister effect, in which ions modulate the strength of the hydrophobic effect—a major driving force in protein folding and association^7,8^— has been known since at least the nineteenth century^9–11^. Biomolecular interactions involving highly charged nucleic acids—such as DNA:protein interactions critical for DNA repair^12^—have been observed to show sensitivity to macroscopic salt concentrations^13^, as have DNA:antibiotic interactions^14^. In the realm of pharmaceutical design, where there is great interest in engineering small molecule ligands, salt effects are known to modulate the interactions of small molecules with proteins^15^ or with supramolecular hosts^16^.

### Current simulation practice arbitrarily fixes microscopic salt composition

In contrast to real physical systems, where the local region near the biomolecule is able to exchange ions with a macroscopic reservoir at a fixed salt concentration (Figure 1, *middle*), simulations of biomolecules typically fix the *number* of salt molecules present in the simulation volume. There is a great deal of diversity in how the fixed number of added ions is typically determined: Along with the specified macroscopic ion concentration, simulation packages may make use of the total cell volume (e.g., Gromacs^17^), the total solvent volume excluding the biomolecular solutes (e.g., CHARMM-GUI^18^), or the number of water molecules (converting the ion concentration into mole or mass fraction, as in OpenMM^19,20^). Some simulation packages choose to use only minimal neutralizing counterions or no counterions at all, relying on uniform background neutralizing charge to allow treatment of long-range electrostatics by particle mesh Ewald (PME) methods^21,22^ (such as Schrödinger’s FEP+ alchemical free energy calculations^23^). In simulation volumes large enough to mimic the inclusion of a macroscopic salt reservoir far from the biomolecular system of interest, the environment near the biomolecule may be accurately represented, but long correlation times for well-ordered ions may still hinder equilibration of the ion environment^24–26^.

### Simulations in the semigrand canonical ensemble can mimic real salt fluctuations

Simulations in the (semi)grand canonical ensemble, however, can—at least in principle—remedy this situation by explicitly allowing one or more components (such as ions) to fluctuate over the course of the simulation via *grand canonical Monte Carlo* (GCMC) moves (Figure 1, *right*). In grand and semigrand canonical methods, simulations are placed in thermodynamic equilibrium with a theoretical reservoir of components. The simulation can exchange molecules/particles with the reservoir, and the concentration the components in the reservoir are specified by their respective chemical potentials. Before running these simulations, one first has to determine the mapping between the concentration in the reservoir and chemical potentials, a process we refer to as *calibration*. Sampling over ion concentrations in explicit water via straightforward GCMC is difficult: Monte Carlo insertion/deletions have to overcome long-range effects, low acceptance rates for instantaneous Monte Carlo moves, and the concentration is sensitive to small (< *k*_*B*_*T*) variations in the chemical potential. Some efforts have circumvented these issues by using implicit solvent models^6,27^, cavity-biased insertions in specialized solvent models^28^, and explicit solvent reorganization moves^29^. *Osmotic ensemble Monte Carlo* schemes that use fractional ions and Wang-Landau approaches have also proven themselves to be useful in simulations of simple aqueous electrolytes^30,31^.

### Nonequilibrium candidate Monte Carlo (NCMC) can achieve high acceptance rates

More recently, nonequilibrium candidate Monte Carlo (NCMC) has been shown to be an effective solution to the problem of low acceptance rates when inserting or deleting particles^32^. In contrast to an instantaneous Monte Carlo (MC) proposal in which an inserted particle is switched instantaneously on and may clash with other solvent or solute particles, in an NCMC proposal, the particle is switched on slowly as the system is allowed to relax via some form of dynamics. NCMC uses a modified acceptance criteria that incorporates the nonequilibrium work to ensure that the resulting endpoints sample from the equilibrium distribution. With well-tuned nonequilibrium protocols, NCMC acceptance rates can be astronomically higher than their instantaneous MC counterparts^32^. In work simulating biomolecules at constant-pH, for example, Roux and coworkers have demonstrated how NCMC is effective at achieving high acceptance rates for NCMC proposals that also transmute an ion to/from a water molecule to maintain net charge neutrality of the system^33,34^.

While calibration of the effective chemical potential for the water and ion forcefields and simulation parameters at hand is nontrivial, this technical challenge can be satisfyingly addressed with existing technologies: Self-adjusted mixture sampling (SAMS)^35^, a form of adaptive expanded ensemble sampling^36^, can be used to conveniently achieve uniform sampling of all relevant salt concentrations in a single simulation, while the Bennett acceptance ratio (BAR) can optimally extract estimates of the relevant free energy differences from all NCMC proposals along with good estimates of statistical error and minimal bias^37–39^. Independent simulations at each salt concentration could be performed separately, with nonequilibrium switching trajectories used to estimate relative free energies between different numbers of salt pairs. However, SAMS helps more rapidly decorrelate the configurations of ions and, in principle, allows a single simulation to be used for calibration.

### An NCMC osmostat can be used alongside thermostats and barostats

Here, we present a new approach that makes use of NCMC to insert/delete salt pairs with high acceptance probability in a manner that correctly models the statistical mechanics of exchange with a macroscopic salt reservoir. The osmostat needs to be calibrated once for the specified solvent and ion models, simulation parameters, and thermodynamic conditions (temperature, pressure, pH, etc.). Following calibration, the osmostat is used in a manner similar to a Monte Carlo barostat, attempting to modify the system composition (and hence interaction potential) at regular intervals to ensure sampling from a target probability density that models a system in equilibrium with a macroscopic salt reservoir (Figure 2). Similar to a Monte Carlo barostat^19,40^, the osmostat moves can be integrated alongside molecular dynamics simulations and other Monte Carlo schemes to sample from equilibrium distributions with specified thermodynamic control parameters. This composability is a general feature of Markov chain Monte Carlo moves, which provide a useful framework for designing modular algorithms for biomolecular simulation^41^.

**Figure 2.**
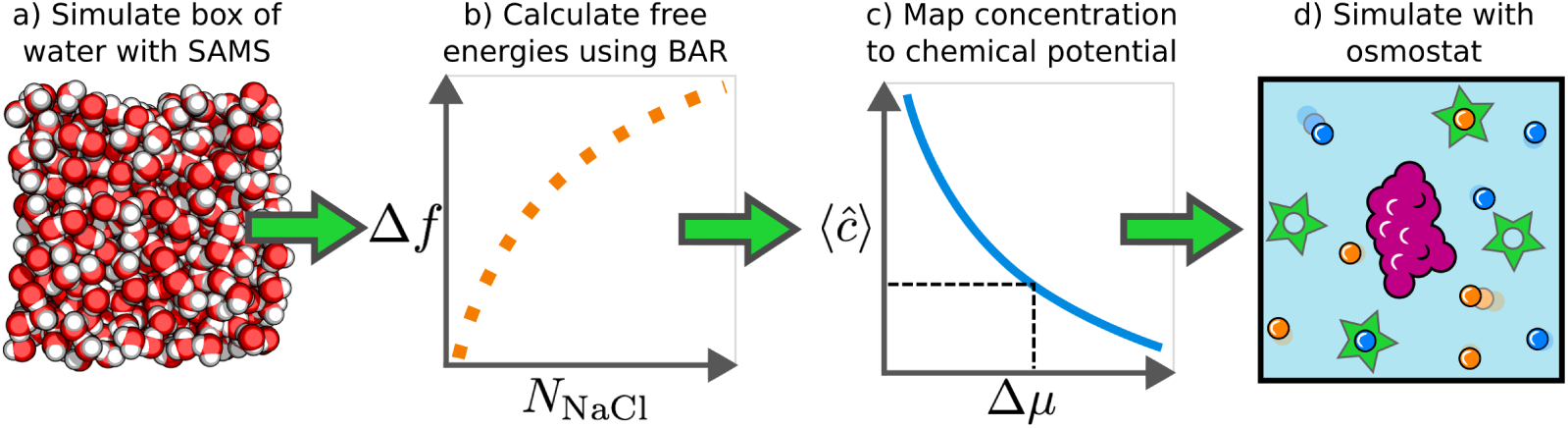
Schematic illustration of the workflow used to calibrate and implement the osmostat. *(a)* Self-adjusted mixture sampling (SAMS) simulations sample an entire range of salt pairs, *N*_NaCl_ ∈ [0, *N*_NaCl, max_], in a sufficiently large box of water to model a saline reservoir. Nonequilibrium candidate Monte Carlo (NCMC) is used to achieve high acceptance rates during salt insertion/deletion attempts, in which an NaCl molecule is transformed into a pair of water molecules, or vice versa. *(b)* The Bennett acceptance ratio (BAR) estimator uses the work values from *all* NCMC proposals (including rejected proposals) to compute an optimal estimate of the (dimensionless) relative free energy, Δ*f*(*N*_NaCl_) ≡*f*(*N*_NaCl_ + 1)- *f*(*N*_NaCl_), to add an additional NaCl salt pair to the box of saline as a function of the number of salt pairs already present, *N*_NaCl_. BAR allows *f*(*N*_NaCl_) to be estimated to a higher precision than the estimates from SAMS. *(c)* Once Δ*f*(*N*_NaCl_) has been computed for the desired water/ion forcefield and simulation parameters governing the energy computation (such as long-range electrostatics treatment), the chemical potential Δ*µ* that produces the desired macroscopic salt concentration ⟨*ĉ*⟩ is numerically computed using equation 19. *(d)* This same chemical potential Δ*µ* is subsequently used as the thermodynamic parameter governing the osmostat to simulate a biomolecular system in equilibrium with an infinitely sized saline reservoir at the specified macroscopic salt concentration.

### How do salt environments vary in realistic biomolecular simulations?

Once we have developed and validated this tool, we use it to ask biophysical questions about the nature of salt environments around biological macromolecules: What is the average salt concentration in the simulation volume, and how does it compare to bulk? Which heuristic scheme, if any, most closely approximates the local salt concentration: macroscopic concentration times total cell volume or solvent volume, or mole fraction of water molecules? How much does the local salt concentration and ionic strength vary in “typical” biomolecular simulation conditions for different classes of biomolecular systems, such as proteins and nucleic acids? And can a Monte Carlo osmostat reduce correlation times for ions over that seen in standard MD simulations, such as the slow correlation times in ion environments around nucleic acids^25^? We consider some test systems that represent different classes of common biomolecular simulations: TIP3P^42^ (and TIP4P-Ew^43^) water boxes, dihydrofolate reductase (DHFR), the *apo* kinase Src, and the Drew-Dickerson B-DNA dodecamer^25^ as a typical nucleic acid.

### Outline

This paper is organized as follows: First, we review the theory behind (semi)grand canonical ensembles that model the fluctuations experienced by small subvolumes surrounding biomolecules. Second, we describe the algorithmic design of the osmostat used to allow salt concentrations to fluctuate dynamically. Finally, we apply the osmostat to address biophysical questions of interest and discuss the nature of salt distributions and their fluctuations.

### Theory and methodology

#### An NCMC osmostat for sampling ion fluctuations in the semigrand ensemble

An *osmostat* is like a thermostat or barostat but allows the number of salt pairs in the simulation box to change dynamically under the control of a conjugate thermodynamic parameter—here, the chemical potential of salt. Salt pairs can be thought of as being exchanged with a macroscopic reservoir, with the free energy to add or remove salt to this reservoir described by the applied chemical potential. In principle, an osmostat could be implemented by including a number of noninteracting (“ghost”) molecules in the simulation volume, turning their interactions on and off to allow the number of active salt molecules to fluctuate dynamically; alternatively, new salt molecules could be introduced or removed dynamically using reversible-jump Monte Carlo (RJMC) methods^44^. In either case, solvent cavity formation to accommodate ions would almost certainly require nonequilibrium protocols that employ soft-core potentials and significant tuning of these insertion/deletion protocols to achieve high acceptance rates.

To simplify implementation for the ions most commonly used in biomolecular simulations (such as NaCl or KCl), we instead choose to exchange the *identities* of water molecules and salt ions, where our conjugate thermodynamic parameter 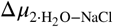 (which we will abbreviate as Δ*µ*) will represent the difference in chemical potential between withdrawing an NaCl molecule from the reservoir while returning two H_2_O molecules. Because solvent cavities are not being created or destroyed—only modified slightly in size—this should provide superior phase space overlap between initial and final states.

We denote the total number of water molecules and ions as *N*, and define the identities of the water molecules and ions with the vector *θ* = (*θ*_1_, *θ*_2_, …, *θ*_*N*_) with *θ*_*i*_ ∈ {−1, 0, +1} to denote anions (*θ*_*i*_ = −1), water (*θ*_*i*_ = 0), and cations (*θ*_*i*_ = +1), respectively (with the potential to extend this to divalent ions by adding -2, +2).

This choice of labeling allows us to define the total number of Na^+^ ions as

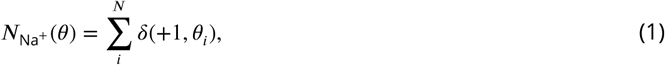

the total number of Cl- ions as

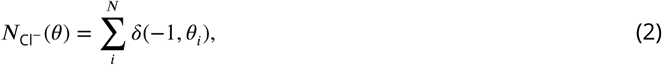

and the number of water molecules as

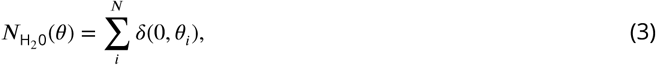

where *δ*(*x, y*) denotes the Kronecker delta, which is unity when *x* = *y* and zero otherwise, and sums run from *i* to *N*. Note that the total number of waters and ions, 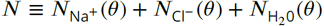, is fixed, and does not depend on *θ*. We define the total charge number of the biomolecules, excluding counterions, as *z*.

When *z* ≠ 0, counterions will be added to ensure that the total charge of the simulation system is zero. The system can be neutralized by any of choice of *θ* that satisfies *n*(*θ*) = −*z*, where the total charge due to ions is given by

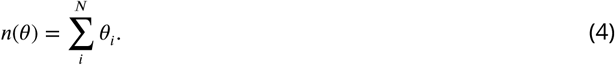

As neutralizing the system will lead to unequal numbers of Na^+^ and Cl^−^, we define the amount of salt as the number of neutral pairs,

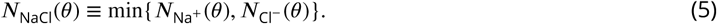

### The semigrand ensemble models salt exchange with a macroscopic salt reservoir

When our osmostat is combined with a scheme that samples the isothermal-isobaric (*N, p, T*) ensemble, we formally sample the semigrand-isothermal-isobaric ensemble (Δ*µ, N, p, T*). The associated equilibrium probability density is given by

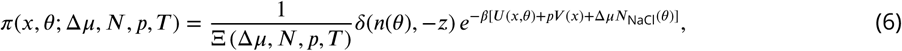

where the Kronecker delta *δ*(*n*(*θ*), −*z*) imposes net charge neutrality, *β* ≡ 1/*k*_*B*_*T* is the inverse temperature, and Ξ (Δ*µ, N, p, T*) is the normalizing constant, given by

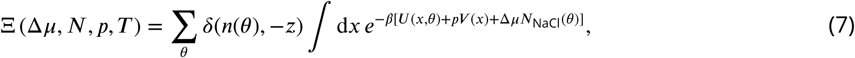

where the outer sum is over all identity vectors and the integral is over all configuration space. For brevity, the dependence of *π* and Ξ on *z* will be omitted. It is also possible to express the probability density of the system as a function of the total number of cations and anions, rather than as function of *θ*. This can be achieved by summing *π*(*x, θ*; Δ*µ, N, p, T*) over all identity vectors that preserve the neutral charge of the system and *N*_NaCl_(*θ*) at some constant value *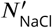*:

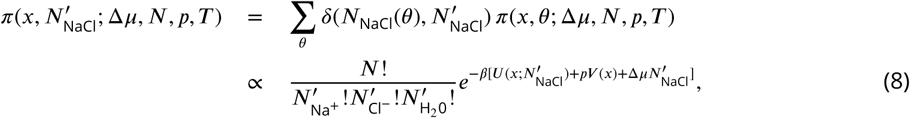

where 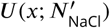 is the potential energy for a system with fixed particle identities that contains 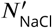 salt pairs. The factorial prefactors account for the degeneracy number of identity vectors *θ* that satisfy the constraints 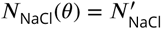 and *n*(*θ*) + *z* = 0.

### Gibbs sampling provides a modular way to sample from the semigrand ensemble

A Gibbs sampling framework can be used to create a modular simulation scheme in which the osmostat updates molecular identities infrequently while some MCMC scheme (such as Metropolis Monte Carlo or Metropolized molecular dynamics) updates particle positions using fixed particle identities:

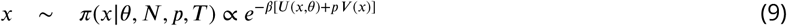

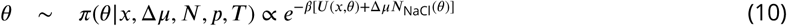

By embedding this approach in a Gibbs sampling framework, it allows the osmostat to readily be combined with other sampling schemes that make use of a Gibbs sampling framework such as replica exchange and expanded ensemble simulations^45^.

Instead of instantaneous MC switching to propose changes in the chemical identities *θ* at fixed configuration *x*, nonequilibrium candidate Monte Carlo (NCMC) is used to propose updates of chemical identities and positions simultaneously as sufficiently long switching trajectories can sampling efficiencies that are orders of magnitude larger than instantaneous proposals^32^:

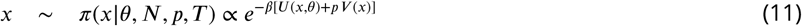

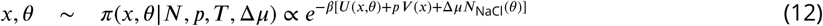

NCMC uses a modified Metropolis-Hastings acceptance protocol in which the appropriate *total work* for switching is accumulated during the nonequilibrium proposal and used in the acceptance criterion.

### The chemical potential Δ*µ* must be calibrated to model macroscopic salt concentrations

Simulating a system that is in chemical equilibrium with an infinitely large saline reservoir at a specified salt concentration first requires the calibration of the chemical potential Δ*µ*. There are multiple ways that one could compute the necessary chemical potential. For instance, one could approximate the reservoir with a sufficiently large box of water, and narrow-in on the chemical potential that produces the desired salt concentration using stochastic approximation or the density control method recommended by Speidal et al.^46^. However, this requires carrying out separate calibration calculations for each desired macroscopic concentration. Instead, we aim to construct a simple calibration procedure by computing the free energies to insert salt pairs into a sufficiently large box of water. We then use these free energies to analytically compute macroscopic salt concentrations over a wide range of chemical potentials, providing a relationship that can be numerically inverted. This procedure need be done only once for a specified ion and water model, though it may need to be repeated if the method used to compute long-range electrostatic interactions is modified.

Our calibration method is similar in principle to that of Benavides et al.^47^, who estimated the chemical potential of NaCl by calculating the free energy to insert NaCl to over a range of concentrations. However, unlike^47^—where the goal was to estimate the solubility of NaCl—our interest in estimating the chemical potential lies solely in its ability to determine the chemical potential of the osmostat saline reservoir corresponding to the desired macroscopic salt concentration in order to induce the appropriate salt distribution on microscopic simulation systems.

Our approach to calibration computes the free energies to add *N*_NaCl_ ∈ {1, 2, …, *N*_NaCl, max_} salt pairs to an initially pure box of water. We limit our free energies calculations to insert NaCl up to some maximum *N*_NaCl, max_ ≪ *N* for practical convenience. No constraint is placed on the amount of salt that can be added in osmostat simulations—instead, the value of *N*_NaCl, max_ impacts the accuracy with which the osmostat can reproduce high macroscopic salt concentrations. We define the absolute dimensionless free energy of a system with *N*_NaCl_ salt pairs at pressure *p* and temperature *T* as *f*(*N*_NaCl_, *N, p, T*),

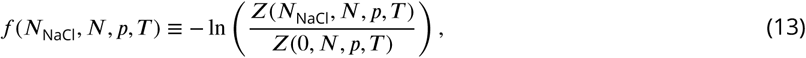

where the partition function 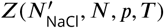 is given by

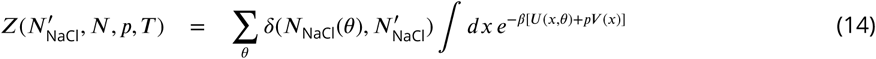

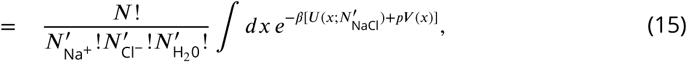

where the number of water molecules 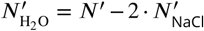. For convenience, we define relative free energies as

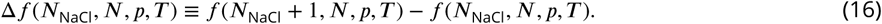

For simplicity, we shall use *f*(*N*_NaCl_) and Δ*f*(*N*_NaCl_) as abbreviations to equations 13 and 16, respectively. The free energies *f*(*N*_NaCl_) can then be used to calculate the average number of salt pairs as a function of the chemical potential Δ*µ*,

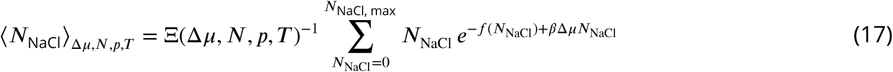

where the semigrand partition function Ξ(Δ*µ, N, p, T*) (the same one from equation 7) can be compactly written as

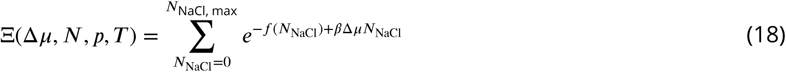

Knowledge of *f*(*N*_NaCl_) will also provide a convenient estimate of the macroscopic salt concentration. We define the macroscopic salt concentration as the mean salt concentration of a system in the thermodynamic limit, and derive in Appendix 2 the following expression for the macroscopic concentration that is amenable to computational analysis:

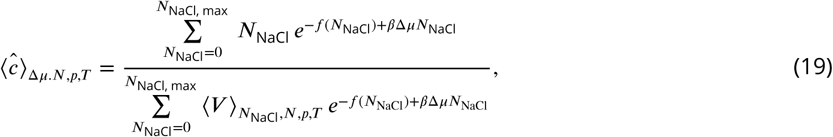

where 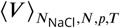 is the average volume for a fixed *N*_NaCl_. The macroscopic concentration ⟨*ĉ*⟩_Δ*µ,N,p,T*_ is a monotonic function of the chemical potential Δ*µ*. Therefore—provided one has estimates of *f*(*N*_NaCl_) and 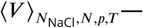 the value of the chemical potential Δ*µ*(*c*) that yields a desired macroscopic concentration ⟨*ĉ*⟩_Δ*µ,N,p,T*_ can be obtained by numerically inverting equation 19.

### Free energies for salt insertion can be efficiently computed using SAMS

One could estimate the free energies *f*(*N*_NaCl_) *N*_NaCl_ ∈ {0, 1, …, *N*_NaCl, max_} using a *N*_NaCl, max_ − 1 equilibrium calculations of the relative free energies Δ*f*(*N*_NaCl_) or the recently developed grand canonical integration technique^48,49^. As the latter requires *a priori* knowledge of the approximate scaling of the chemical potential with the concentration, we instead opt to use the recently proposed self-adjusted mixture sampling (SAMS)^35^ method to facilitate the calculation of the free energies from a single simulation. SAMS is a development on the method of expanded ensembles^36^ (sometimes known as serial tempering^50^) and generalized Wang-Landau algorithms^51,52^. It is a stochastic approximation scheme that produces unbiased estimates of the free energies (unlike Wang-Landau) that—in the asymptotic limit—have the lowest variance out of all other stochastic approximation recursion schemes^35^. It can be used to sample over a discrete state space and simultaneously estimate the relative log-normalizing constant for each state. For our calibration simulations, the discrete states correspond to the number of salt pairs in the systems *N*_NaCl_ ∈ {0, 1, …, *N*_NaCl, max_} and the log-normalizing constant are the desired free energies *f*(*N*_NaCl_). By dynamically altering a series of biasing potentials, one for each state, the SAMS algorithm asymptotically samples the discrete states according to user specified target weights^35^. When the target weights are uniform over the state space—as we choose herein to ensure the uncertainties in the estimated free energies are approximately equal—the biasing potentials are themselves estimates of the free energies *f*(*N*_NaCl_). Thus, SAMS can, in principle, calculate all *f*(*N*_NaCl_) in a single simulation more efficiently and conveniently than numerous independent equilibrium free energy calculations.

As we describe below, our osmostat employs NCMC, which allows us to calculate the salt-insertion free energies by processing all of the NCMC protocol work values in the SAMS simulations with BAR, even from the attempts that are rejected. BAR requires samples of forward and reverse work samples of salt insertion and deletion attempts to compute Δ*f*(*N*_NaCl_) and its statistical uncertainty for *N*_NaCl_ ∈ {0, 1, …, *N*_NaCl, max_}^37–39^. These relative free energies can then be summed to estimate *f*(*N*_NaCl_) and corresponding statistical uncertainties. Our calibration simulations therefore exploit the sampling efficiency of SAMS and the estimation efficiency of BAR.

In general, the chemical potential Δ*µ* will need to be recalibrated if the practitioner changes temperature, pressure, water or ion forcefield models, nonbonded treatment, or anything that will affect *f*(*N*_NaCl_) or 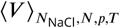. A sufficiently large water box must be used when calculating *f* (*N*_NaCl_) to reach a regime in which *f*(*N*_NaCl_) is insensitive to changes in simulation size; as we will show, our calibration simulations achieve this size insensitivity even for modest water boxes of a few thousand molecules.

### The osmostat maintains electrostatic neutrality

To use PME^21^, a popular choice for accurate long-range electrostatics, charge neutrality of the entire system needs to be maintained to avoid the artifacts induced by application of a uniform background neutralizing charge^22^. Even if an alternative long-range electrostatics treatment is employed (e.g. reaction field electrostatics or other non-Ewald methods^53^), there is, in general, approximate equality between the total number of negative charges and positive charges in biological microenvironments as they approach macroscopic lengthscales (see Figure 1 *left*). From a purely theoretical perspective, the existence of a thermodynamic limit a system with a net charge depends on the particular details of the system^54^. For these reasons, we ensure that our proposals always maintain charge neutrality by inserting or deleting a neutral Na^+^ and Cl^−^ pair.

We insert and delete a salt pair by converting Na^+^ and Cl^−^ ions to two water molecules (see Figure 3). These moves convert the nonbonded forcefield parameters (partial changes *q*, Lennard-Jones radii *σ*, and Lennard-Jones potential well-depths *ϵ*) of the water and ion parameters. The Na^+^ and Cl^−^ ions are given the same topology, geometry, and number of atoms as the water model used for the simulation. Irrespective of the choice of water model, the nonbonded ion parameters are placed on the water oxygen atom, and the hydrogen atoms or additional charge sites (such as in TIP4P) have their nonbonded interactions switched off. The manner in which salt and water are transmuted to one another are is described in Appendix 3. The mass of the ions is set as the same as water, which has no impact on the equilibrium configuration probability density, though it may distrupt the kinetics (which are not of interest here).

**Figure 3.**
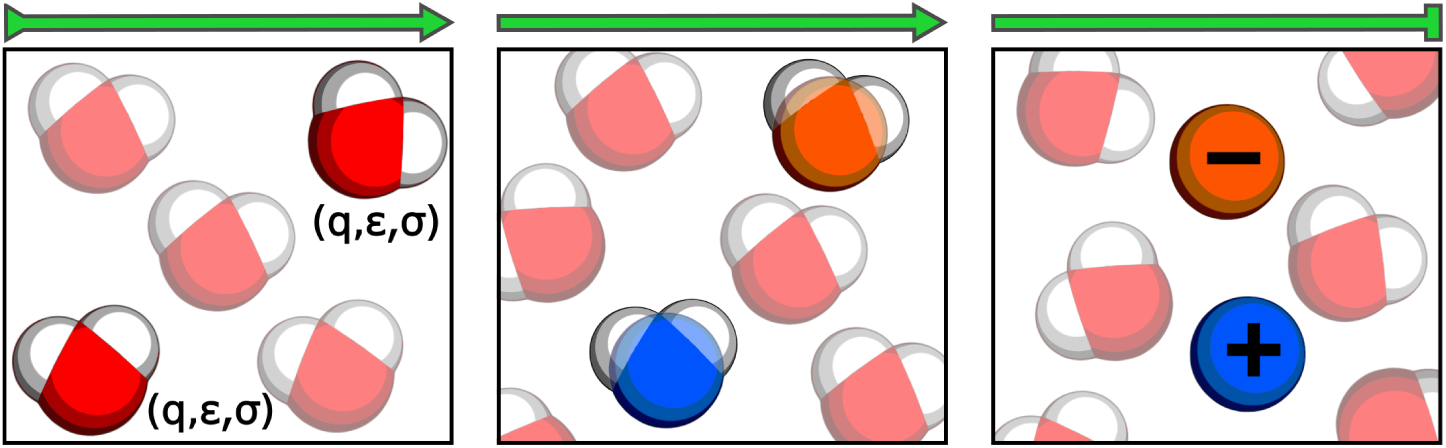
Schematic illustration of the nonequilibrium candidate Monte Carlo (NCMC) alchemical protocol used to insert NaCl. Two water molecules are chosen at random for transformation into Na^+^ (blue sphere) and Cl^−^ (orange sphere). Over a number of NCMC steps, the nonbonded parameters of each atom in the water molecules, namely the partial charges, *q*, Lennard-Jones energy well depths, *ϵ*, and Lennard-Jones separation parameters, *σ*, are transformed into the nonbonded parameters of the ions along a linear interpolation of the parameters. The hydrogen atoms and extra charge sites (if present) of the water model remain attached to the ions as non-interacting dummy atoms. The entire NCMC proposal is then accepted or rejected according to the probability given in equation 56. Note that osmostat NCMC moves are mixed with standard Langevin integration at a fixed timestep to obtain fully ergodic sampling. A full description of the Monte Carlo and NCMC procedure used here is provided in Appendix 3.

### Nonequilibrium candidate Monte Carlo is used to enhance sampling efficiency

A benefit of exchanging ion and water nonbonded forcefield parameters is that this procedure avoids the need to create new cavities in solvent, a difficulty that significantly complicates particle creation and destruction techniques. Nevertheless, instantaneous Monte Carlo attempts to interconvert salt and water will be overwhelmingly rejected as it is highly unlikely that the dipoles of the molecules that surround a transmuted ion—usually solvent—will be orientated in a manner that favorably solvates the new charge. This effect is compounded by the long-range nature of Coulombic interactions. The acceptance probability for salt insertion and deletion would improve drastically if the dipoles and locations of the solvent could be redistributed during an MCMC attempt. Previously, Shelly and Patey developed a configuration bias Monte Carlo technique for the insertion and deletion of ions in grand canonical Monte Carlo^29^. Their method reorients dipoles in a shell surrounding the inserted or deleted ion, which improved the sampling efficiency by over two orders of magnitude^29^.

Here, we use nonequilibrium candidate Monte Carlo (NCMC)^32^, a technique that is closely related to sequential Monte Carlo and annealed importance sampling^55,56^, to automatically relax systems around inserted or deleted ions, thereby boosting acceptance rates and sampling efficiencies to values far higher than reported elsewhere.

In NCMC, a Monte Carlo attempt is divided into a nonequilibrium protocol that drives the system through many intermediate states. Candidate configurations are generated by driving a chosen set of variables (thermodynamic or configurational) through these intermediate states whilst allowing unperturbed degrees of freedom to relax via dynamical propagation in response to the driving protocol. The total amount of work that is accumulated between interleaved steps of perturbation (of the variables of interest) and propagation (of the unperturbed degrees of freedom) is used to accept or reject the candidate configuration. Good NCMC acceptance rates can be achieved for a reasonable choice of nonequilibrium protocol; often, a parametric protocol is specified and the total protocol length (or *NCMC switching time*) is tuned to be long enough to ensure a system is sufficiently relaxed with respect to the completed perturbation but short enough to be efficient.

In our NCMC osmostat, the nonbonded parameters of the ions and water molecules being exchanged are linearly interpolated into a series of equally spaced alchemical states. Each perturbation step along the alchemical path was followed by a fixed number of time-steps of Langevin dynamics where the configurations of the whole system were integrated (see Figure 3). A full description of our Monte Carlo and NCMC procedure is provided in Appendix 3. Here, NCMC propagator uses the same Langevin integrator as used in equilibrium sampling to ensure there was no significant mismatch between the sampled densities. Our particular choice of Langevin integrator (described below) was used to avoid the long correlation times that results from fully Metropolized molecular dynamics integrators and to mitigate the configuration sampling bias that is incurred by unmetropolized finite time-step integrators.

### We use an integrator that minimizes configuration sampling bias

Care must be taken to ensure that the total work is properly accumulated in NCMC, as incorrect accumulation of work or the use of alternative definitions will lead to erroneous computation of the acceptance probability and simulation results. For time reversible MCMC integrators, such as with generalized Hamiltonian Monte Carlo (GHMC), the total work is the *protocol work*: the sum of the instantaneous potential energy changes that result from each perturbation during the driving process^57^. If the system is relaxed in-between perturbations using propagators that do not leave the target distribution invariant, such as unmetropolized Langevin integrators, NCMC can drive systems to undesirable nonequilibrium steady states, whose statistics may differ from equilibrium. On top of the work that is already performed by the driving protocol, propagators that do not satisfy microscopic reversibility can also be considered to perform work on a system^57^. This work, known as the *shadow work*, must either be minimized or eliminated (i.e., via Metropolizing the dynamics) for NCMC to sample very close to, or exactly, from the target probability density.

The issue of shadow work accumulation is not limited to propagators in NCMC. Indeed, *all* finite time-step molecular dynamics integrators incur a discretization error that results in biased sampling when used without metropolization. While configuration sampling errors do not occur with GHMC, the correct acceptance criterion requires that the momenta of all particles are reversed upon rejection (or acceptance) of a proposal. The reversal of momenta results in a simulation ‘retracing its steps’, thereby significantly increasing correlation times and decreasing sampling efficiencies. Hamiltonian Monte Carlo sampling can suffer from even longer correlation times, as momenta are randomized for each trial, irrespective of whether the previous move was accepted or not. This problem can be mitigated by using GHMC reduced momentum flipping schemes that still rigorously sample from the target distribution^58–60^. Correlation times are minimized by GHMC schemes that do not reverse momenta at all, although this incurs sampling bias^61^.

Recently, Leimkuhler and Matthews have proposed an unmetropolized Langevin dynamics technique that incurs minimal configuration sampling bias^62^. The minimal error is achieved using a particular numerical scheme to update the positions and momenta at each time-step. Denoting half time-step velocity updates as V, half time-step position updates as R, and the addition of an Ornstein-Uhlenbeck process as O (the Brownian motion “kick”), the symmetry in the VRORV splitting scheme leads to a particularly favorable cancellation of configuration sampling error. Leimkuhler and Matthews also found that than VRORV exhibited the lowest error on configuration dependent quantities, such as the potential energy, in biomolecular simulations compared to other symmetric splittings. As Langevin dynamics with VRORV splitting samples very closely to the true configuration Hamiltonian, we expect its neglect within NCMC moves designed to sample configurational properties to induce very little error in sampled configurational densities. For this reason, we used the protocol work to accept or reject proposals from NCMC in our osmostat.

#### Salt concentration and ionic strength

##### Ionic strength influences the effective salt concentration

We are interested in quantifying the variation of the instantaneous salt concentration *c* in our osmostated biomolecular simulations, where

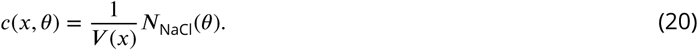

Although the salt concentration of the saline reservoir, i.e. the macroscopic concentration, is known precisely and controlled by the user, the presence of a biomolecule in a simulation, along with any neutralizing counterions, may lead to significant differences in the mean salt concentration in the simulation volume from the macroscopic salt concentration. In contrast, the mean salt concentration in an initially pure box of water should match the macroscopic salt concentration of the reservoir if the chemical potential used in the osmostat is accurately calibrated.

The Debye-Hückel theory of electrolytes provided an early, analytical treatment of dilute ionic solutions using continuum electrostatics. In Debye-Hückel theory, the ionic strength *I* of a system, which for our simulations is

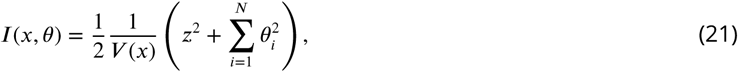

is used to predict how the effective concentrations, or activities, of ions are affected by the presence of electrolytes in the solution. The key insight of Debye-Hückel theory is that—because of electrostatic screening—the ionic strength tempers the activity of ions, such that increasing the ionic strength of a solution lowers the effective concentration of electrolytes. Although Debye-Hückel theory is too simplistic to be used to accurately predict the salt concentration in biomolecular simulations, the ionic strength may still provide insight into the salt concentrations that we will observe in our osmostated simulations. Thus, we will investigate the variation of the ionic strength as well as the salt concentration. As a large charge number of the biomolecule *z* will dominate *I* for small simulation volumes, we will also consider the variation of ionic strength of the solvent only, i.e., by neglecting *z*^2^ in equation 21.

##### Simulation packages add different amounts of salt

There is diversity in the way that current practitioners of all-atom biomolecular simulations add salt (*salinate*) to systems during the preparation stages of simulations. While it is common that only neutralizing counterions are added, a number of workflows elect not to add counterions at all^23^. Salt pairs may be added, or not added at all, and when they are added, simulation packages use differing definitions of salt concentration, such that each package can add different numbers of salt pairs to the same system even if the desired salt concentration is the same. All packages ignore the presence of neutralizing counterions when adding salt. In this study, we are concerned with quantifying the accuracy of some of the most popular salination techniques.

Given a target salt concentration of *c*_*t*_, a popular method to add salt—exemplified by the Gromacs package^17^—uses the initial volume of the system *V* (*x*_0_) to count the required number of pairs. We determine the number of salt pairs that would be added by this strategy as

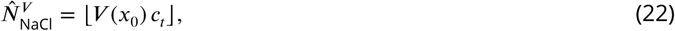

where ⌊*y*⌋ denotes the floored value of *y*. We are interested in assessing the accuracy of the corresponding concentration of salt 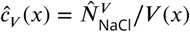. Preparation tools such as CHARMM-GUI^18^ add salt based on the initial volume of the *solvent* 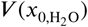, which we reproduce with

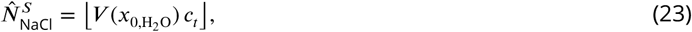

to estimate the corresponding concentration 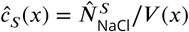 that would occur for all later configurations. Estimates that use strategies similar to equations 22 and 23 are sensitive to initial volume of the system; if salt is added before the volume is sufficiently equilibrated, the salt concentration during the simulation can deviate significantly from the target concentration. In contrast, packages such as OpenMM^19,20^, use the *ratio* of salt pairs to water molecules in bulk solvent to add

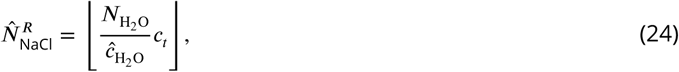

salt pairs, where 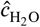 is concentration of bulk water, for which 55.4 M is used by OpenMM. The corresponding salt concentration 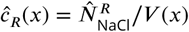, as well as *ĉ*_*V*_ (*x*) and *ĉ*_*s*_(*x*) will be compared to the concentration of salt that results from the application of our osmostat to help inform future simulation strategies.

### Simulation details

#### Systems considered in the study

The primary aims of this study are to quantify and understand how the concentration of salt and ionic strength vary around typical biomolecules, to assess the accuracy of methods that insert salt in typical simulation strategies, and to ascertain whether an NCMC osmostat can decorrelate biomolecule:ion interactions faster than fixed-salt dynamics. To meet these aims, we considered four biological systems that are representative of those that are commonly simulated with molecular dynamics: pure water, dihydrofolate reductase (DHFR), the *apo* kinase Src, and the Drew-Dickerson B-DNA dodecamer palindromic sequence. All systems were taken from the OpenMMTools [0.11.1] set of test systems^63^, such that each system has a different provenance.

Dihydrofolate reductase (DHFR) is a small, globular enzyme that has frequently been used as a model system in molecular simulations. The DHFR structure used here was taken from the joint Amber-CHARMM (JAC) benchmark (obtained from the Amber 14 benchmark archive^64^). The protein structure was stripped of hydrogen atoms, and using `tleap`^65^, was re-protonated at pH 7 and solvated in an orthorhombic box of TIP3P waters that had a clearance of at least 10 Å. The Amber 14SB forcefield from the AmberTools 16 package was used for the protein^65^. As an initial relaxation of the system, the solvated system was minimized and propagated for 3 ps with Langevin dynamics.

The tyrosine kinase Src, a member of the non-receptor tyrosine kinase family, was selected for this study as an example of a prototypical drug target. The *apo* Src structure was taken from the OpenMMtools `testsystems` data set and resolvated with TIP3P in an orthorhombic box that was at least 10 Å away from the protein. As part of the preparation, the energy of system was minimized and subsequently relaxed using 3 ps of Langevin dynamics to remove any bad contacts. Further equilibration was performed as detailed below. The original system was not suitable for simulation with the osmostat as fixed neutralizing counterions were present in the system. The OpenMMtools structure was downloaded from the Protein Data Bank, identification code 1YI6, and prepared using PDBFixer^66^ and protonated at pH 7. The small molecule in the binding site was also removed during the preparation. The Amber 14SB forcefield from the AmberTools 16 package was used for the simulations^65^.

The Drew-Dickerson dodecamer (CGCGAATTGCGC) is a classic model DNA system. The B-DNA structure of the Drew-Dickerson dodecamer was downloaded from the Protein Data Bank (identification code 4C64). The structure was stripped of ions and solvated in a box of TIP3P water to ensure at least 9 Å of clearance around the DNA. To test the effect of the amount of solvent on the distribution of salt and ions, the structure was also solvated in a box of TIP3P water that had a clearance of at least 16 Å around the DNA. As with the *apo* kinase Src, the system was energy minimized and subsequently relaxed using 3 ps of Langevin dynamics. As described below, further equilibration was also performed. The Amber OL15 forcefield from the AmberTools 16 package was used for the DNA^67^.

#### General simulation details

Simulations were performed with OpenMM [7.1.0]^20^. The osmostat was implemented within the open-source package SaltSwap [0.5.2] that was written for the purpose of this publication. Simulations utilized either TIP3P^42^ or TIP4P-Ew^43^ water models, and Joung and Cheatham parameters were used for Na^+^ and Cl^−^ ions^68^. Unless otherwise stated, the amount of salt in a simulation was initialized by salinating the system according to equation 24 with the macroscopic concentration as the target concentration *c*_*t*_.

For all simulations, long-range electrostatic interactions were treated with particle mesh Ewald (PME), with both direct-space PME and Lennard-Jones potentials making use of a 10 Å cutoff; the Lennard-Jones potential was switched to zero at the cutoff over a switch width of 1.5 Å to ensure continuity of potential and forces. PME used a relative error tolerance of 10^−4^ at the cutoff to automatically select the *α* smoothing parameter, and the default algorithm in OpenMM was used to select Fourier grid spacing (which selected a grid spacing of ∼0.8 Å in each dimension). All bonds to hydrogen were constrained to a within a fractional error of 1 × 10^−8^ of the bond distances using CCMA^69,70^, and waters were rigidly constrained with SETTLE^71^. OpenMM’s long-range analytical dispersion correction was used to avoid pressure artifacts from truncation of the Lennard-Jones potential. Simulations were run at 300 K with a Monte Carlo barostat with 1 atm external pressure and Monte Carlo update interval of 25 steps. Equilibrium and NCMC dynamics were propagated using high-quality Langevin integrators taken from the OpenMMTools [0.11.1] package, with a 2 fs timestep and collision rate of 1 ps^−1^. Integrators used deterministic forces and OpenMM’s mixed single and double precision implementation. In addition to the dynamics used to prepare the systems, every simulation was briefly thermalized using 4 ps of dynamics. Where stated, additional simulation data was discarded from the start of simulations using the automatic procedure in the pymbar timeseries module as detailed in^72^. As described above, positions and velocities were updated using the VRORV splitting scheme (also known as BAOAB) to mitigate the configuration space error in equilibrium sampling and NCMC proposals that result from unmetropolized Langevin dynamics^62^

The insertion or deletion of salt was attempted every 4 ps using the procedure described in Appendix 3. All ions used the same number of atoms, topology, and geometry as the water model used in the simulation. As illustrated in Figure 3, the “insertion” of an ion was achieved by switching the nonbonded parameters of the water oxygen atom to either Na^+^ or Cl^−^ and by simultaneously switching the nonbonded parameters of the water hydrogen atoms (along with any extra charge sites) to zero—the “deletion” of an ion involved the reverse procedure. With the exception of the simulations where the NCMC protocol was optimized, the NCMC protocol was 20 ps long, and consisted of 1000 perturbation steps, where each perturbation followed by 10 steps of Langevin integration with a 2 fs timestep. The pseudo-code for the entire NCMC osmostat, including how it is combined with molecular dynamics can also be found in Appendix 3. Unless otherwise stated, the NCMC protocol length is not accounted for in the reported lengths of the simulations.

The simulations were analyzed with open source scripts that used a combination of numpy 1.13.1^73^, scipy 0.19.1^74^, pymbar 3.0.1^75^, MDTraj 1.8.0^76^, VMD 1.9.4^77^ (see *Code and data availability*); the saltswap conda package provided automatically installs the dependencies needed to run the simulation scripts. Plots and figures were produced using Matplotlib 2.0.2^78^ and Inkscape 0.91.

#### Calibration of the chemical potential

The chemical potential was calibrated in cubic boxes of TIP3P water and TIP4P-Ew water. Both boxes initially had edge lengths of 30 Å with water molecules at roughly the same density as bulk water; the box of TIP3P water contained 887 molecules and the box of TIP4P-Ew water contained 886 molecules. Ten 80 ns SAMS simulations were performed on each box, and were targeted to sample uniformly over salt pairs *N*_NaCl_(*θ*) ∈ {0, 1, …, 20}. The insertion or deletion of salt was attempted every 4 ps. Half of the simulations were initialized with 0 salt pairs, whereas the other half were initialized with 20 salt pairs. The maximum number of salt pairs *N*_NaCl, max_ was chosen to be 20 in these calibration simulations because the corresponding salt concentration (roughly 1.2 M) is beyond the concentrations in biological microenvironments that are typically considered. (Note that the maximum amount of 20 salt pairs applies only to these calibration simulations—the osmostat simulations with solutes have no such maximum number of salt pair limitation.) The volumes of the boxes at each salt occupancy were recorded during the SAMS simulations in order to estimate 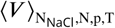 (henceforth abbreviated as 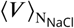. The SAMS simulation procedure automatically provides on-line estimates of the free energies *f*(*N*_NaCl_), which, along 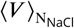, are required to calibrate the chemical potential. The protocol work from all of the NCMC insertion and deletion attempts were post-processed with BAR (using the pymbar package^75^) to provide additional estimates of *f*(*N*_NaCl_) along with statistical uncertainties.

To assess whether *f*(*N*_NaCl_) and 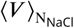 had been accurately calculated, larger boxes of TIP3P and TIP4P-Ew water were simulated for 32 ns at a range of chemical potentials Δ*µ*. The mean salt concentrations from the simulations were compared to concentrations predicted using equation 19 with the estimated values for *f*(*N*_NaCl_) and 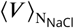. The boxes of these validation simulations were initially 50 Å in length, and contained 4085 TIP3P and 4066 TIP4P-Ew water molecules. These simulations were initialized without any salt present in the systems.

#### Optimization of the NCMC protocol

We consider only two parameters in optimizing the nonequilibrium protocol used in NCMC proposals: the total number of times the potential is perturbed, *T*, and the number of Langevin steps that occur before and after each perturbation, *K*. Generally, we expect the acceptance probability to increase as the overall perturbation is broken into smaller pieces—as *T* increases. Increasing the number of propagation steps following each perturbation, *K*, also improves the acceptance probability in a manner that is dependent on the computational efficiency details of the simulation code. To quantify the trade-off between acceptance probability and compute time, we define the NCMC efficiency *E*(*T, K*) as

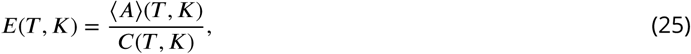

where ⟨*A*⟩ (*T, K*) is the average acceptance probability and *C*(*T, K*) is the average computer time per insertion/deletion attempt. All simulations were performed and timed on single Nvidia GTX-1080 GPUs. The total protocol length of an NCMC attempt is equal to *T* × *K* multiplied by the Langevin integration timestep, which is 2 fs in this case.

Simulations using various NCMC protocols lengths were performed on cubic boxes of TIP3P and TIP4P-Ew that had initial edge lengths of 30 Å. The simulations sampled configurations for a total of 32 ns (excluding the NCMC sampling) and had NCMC protocol lengths up to 40 ps for different combinations of total perturbation steps *T* and propagation steps *K*. The insertion or deletion of salt was attempted every 4 ps, such that there were a total of 8000 insertion/deletion attempts for each simulation. The efficiency of each protocol *E* was estimated relative the efficiency of instantaneous insertion and deletion. Shelly and Patey also used the ratio of the average acceptance probability to the compute time to estimate the efficiency of their configuration bias ion insertion scheme relative to instantaneous insertions^29^. In this work, no effort was made to optimize the alchemical path.

#### Quantifying the scaling behavior of the osmostat

To investigate the sampling efficiency of our osmostat under physiological conditions, DHFR was simulated with macroscopic concentrations of 100 mM, 150 mM, and 200 mM. Each simulation was 30 ns long and there were three repeats per macroscopic concentration. Equation 24 was used to add an initial amount of salt to the simulation. The `timeseries` module in pymbar^75^ was used to estimate the autocorrelation function of salt concentration as well as the integrated autocorrelation time for each macroscopic salt concentration.

It is important to establish how the distributions of salt concentration and salt numbers scale with the number of water molecules in the system and the macroscopic concentration. To this end, we simulated different sizes of water boxes with macroscopic concentrations of 100 mM, 150 mM, and 200 mM. Each simulation was repeated three times.

#### Estimating the efficiency of ion configuration sampling with NCMC

Ponomarev et al. previously used the Drew-Dickerson DNA palindromic sequence to quantify the rate of convergence of spatial ion distributions in DNA simulations^25^. Three osmostated simulations and three fixed-salt simulations of the Drew-Dickerson dodecamer were performed for 60 ns with a macroscopic salt concentration of 200 mM. As the insertion or deletion of salt was attempted every 4 ps, there was a total of 15,000 attempts. The fixed salt simulations used the same ion topologies and masses as those used by the osmostat, are were added to the system using the scheme summarized by equation 24. The autocorrelation of ion:phosphate interaction occupancies were estimated from the osmostated and fixed-salt simulations using the open-source analysis scripts that accompany this manuscript.

#### Quantifying the salt concentration around biomolecules

Three 30 ns simulations of *apo* Src kinase were performed, with salt insertion or deletion attempted every 4 ps, using a macroscopic concentration of 200 mM. The amount of salt that was initially added to this system was calculated using equation 24. These simulations, as well as those of TIP3P water, DHFR, and the DNA dodecamer described above, were used to analyze the distributions of salt concentration (equation 20), ionic strength (equation 21), and the concentrations of salt that would occur for the heuristic salination schemes described in equations 22, 23, and 24.

To further understand the scaling behavior of the distributions of salt concentration with system size, and to assess the extent of finite size effects on the ion spatial distributions around DNA, additional simulations were performed on DNA. The Drew-Dickerson DNA dodecamer was resolvated in a box of TIP3P water that was at least 16 Å away from the molecule. Three repeats of 45 ns long osmostated and fixed-salt simulations were performed, with the insertion or deletion of salt was attempted every 4 ps. The salt concentration distribution was estimated, as were the Na^+^ and Cl^−^ spatial distributions around the DNA.

## Results

### SAMS simulations and BAR estimates accurately capture salt insertion free energies

In order to estimate the chemical potential Δ*µ* corresponding to a desired macroscopic salt concentration, we must have precise estimates of both free energies to insert salt into a box of water containing *N*_NaCl_ salt molecules, *f*(*N*_NaCl_), and the average saline box volume as a function of 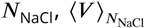, for *N*_NaCl_ ∈ {0, 1, …, *N*_NaCl, max_}. Figure 4 (upper left) depicts the computed relative free energy difference for inserting an additional salt pair into a box of water molecules already containing *N*_NaCl_ salt molecules for both TIP3P and TIP4P-Ew for *N*_NaCl_ ∈ {0, …, 19}. The relative free energies were estimated with BAR using all nonequilibrium work values for salt pair insertion/deletion NCMC proposals, irrespective of whether the proposal attempt was accepted or not, from ten SAMS simulation. Although SAMS also provides *online* estimates for *f*(*N*_NaCl_) over this same range^35^, these online estimates were found to have significantly higher variance than the BAR estimates (see Figure A5.1), so we make use of BAR-derived estimates of *f*(*N*_NaCl_) derived from SAMS simulations throughout.

**Figure 4.**
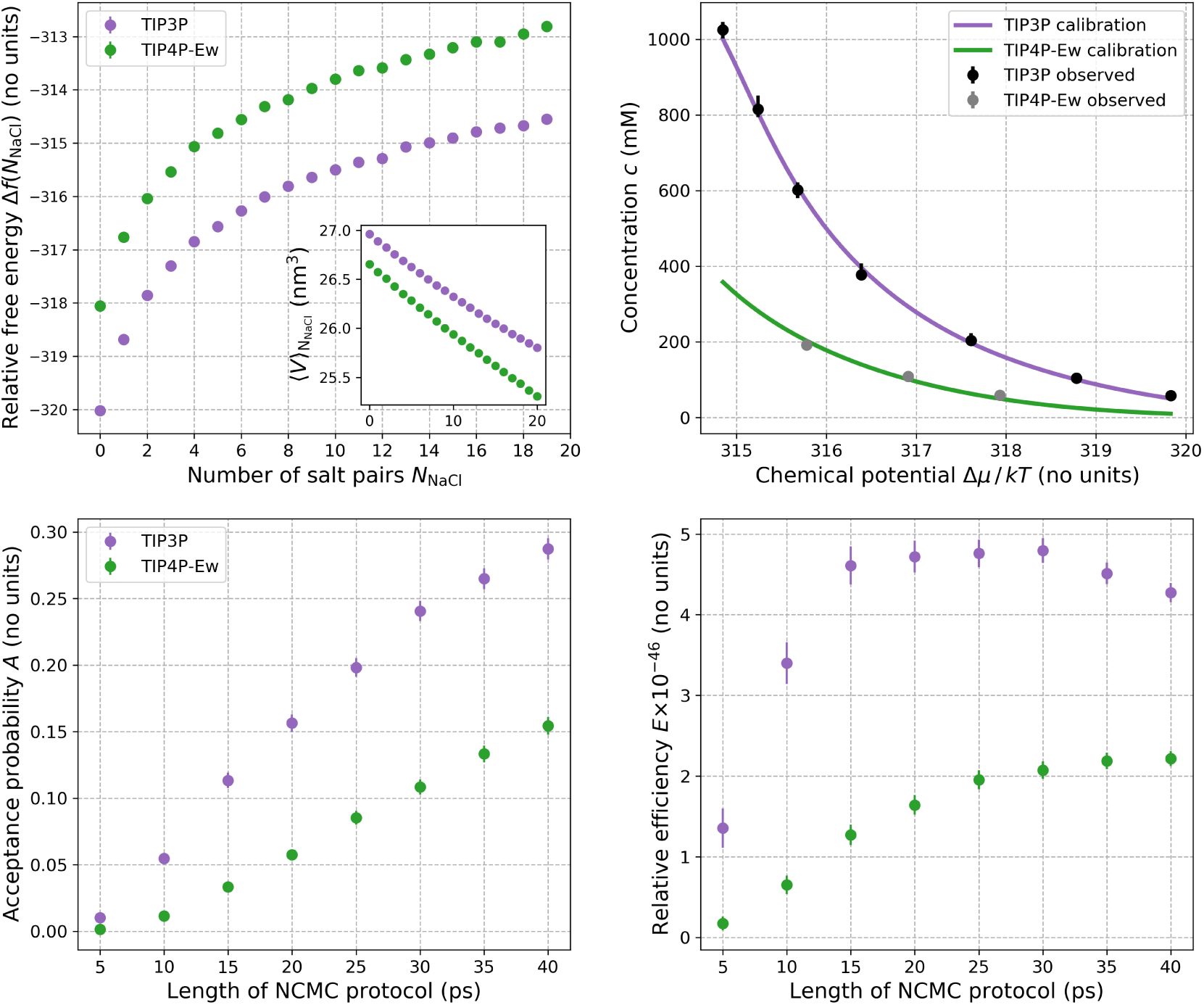
Calibration of chemical potential Δ*µ* for two different water models (TIP3P and TIP4P-Ew) and NCMC protocol optimization. *Top left, main:* The relative free energy Δ*f*(*N*_NaCl_)—estimated from the SAMS calibration simulations— to insert an Na^+^ and Cl^−^ salt pair and remove two water molecules in boxes of TIP3P and TIP4P-Ew water as a function of the number of salt pairs *N*_NaCl_ already present in the box (see equation 16). *Top left, inset:* The average volume 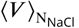 of the saline box as a function of *N*_NaCl_, estimated from the SAMS calibration simulations. The TIP3P box contained a total of 887 molecules (including water and ions) and the TIP4P-Ew box contained 886 molecules. The relative free energies and 95% confidence intervals have been calculated using BAR and are smaller than the circular markers. *Top right:* Predicted relationship between the macroscopic salt concentration ⟨*ĉ*⟩ and chemical potential difference Δ*µ* estimated with equation 19 for TIP3P and TIP4P-Ew (dark lines) compared to the average concentrations ⟨*c*⟩ estimated from equilibrium osmostat simulations of boxes of water at specified chemical potentials (circles). There were 4085 and 4066 molecules in the boxes of TIP3P and TIP4P-Ew water, respectively. Bootstrapping of BAR uncertainty estimates of *f*(*N*_NaCl_) and bootstrap uncertainties of 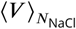 were used to calculate 95% confidence intervals for the mean concentration curves—these fall inside the thick lines. Error bars on the average simulation concentrations show 95% confidence intervals, and have been estimated using bootstrap sampling of statistically independent subsamples of the simulation concentrations. For the osmostat simulations, equilibration times were automatically estimated and independent samples extracted using the timeseries module of pymbar^75^. For these osmostat simulations, the shortest and largest estimated equilibration times were 0.2 ns and 26.9 ns respectively, with the largest equilibration time occurring for TIP3P simulation at the lowest Δ*µ*—the staring salt concentration for this simulation was furthest from the equilibrium value. *Bottom left:* Average acceptance probability for salt insertion and deletion as a function of the NCMC protocol length. Simulations were run with a 200 mM osmostat in boxes of TIP3P (887 molecules) and TIP4P-Ew (886 molecules). The mean instantaneous MC acceptance probabilities for TIP3P and TIP4P-Ew are very small: 3.0 × 10^−51^ [5.0 × 10^−66^, 9.0 × 10^−51^] and 1.0 × 10^−46^ [3.0 × 10^−64^, 4.0 × 10^−46^] respectively, (with 95% confidence intervals denoted in brackets). *Bottom right:* The efficiency (defined by equation 25) of the NCMC protocols relative to instantaneous insertion and deletion attempts in TIP3P for a 200 mM osmostat; all protocols are at least 10^45^ times more efficient than instantaneous insertion and deletion. NCMC protocols of about 20 ps for TIP3P are optimal for our nonequilibrium procedure, though longer protocols are required to achieve similar efficiencies for TIP4P-Ew.

The primary accuracy of the calibration simulations lies in their ability to reproduced desired salt concentrations in bulk water. Nevertheless, it is encouraging to note that calculated free energy to insert one NaCl pair in a box of TIP3P and TIP4P-Ew are broadly in agreement with previous computational estimates and experimental measurements. As implied by equation 16, the free energy to insert the first salt pair, Δ*f*(*N*_NaCl_ = 0), can be expressed as the difference in hydration free energy between NaCl and two water molecules. Assuming the hydration free energy of TIP3P and TIP4P-Ew water to be -6.3 kcal/mol^79^, we estimate the hydration free energy of NaCl to be −171.73 ± 0.04 kcal/mol and −170.60 ± 0.04 kcal/mol in TIP3P and TIP4P-Ew water, respectively. Using a different treatment of long-rang electrostatics but same ion parameters as this study, Joung and Cheatham calculated the individual hydration free energies of Na^+^ and Cl^−^ in TIP3P and TIP4P-Ew, which can be summed to approximate the hydration free energy of NaCl^68^. These hydration free energies (−178.3 kcal/mol in TIP3P -177.7 kcal/mol in TIP4P-Ew) are within 5% of our estimates. For comparison, estimates of standard NaCl hydration free energies based on experimental data are -170.4 kcal/mol^80^, -171.8 kcal/mol^81^, and -177.8 kcal/mol^82^.

### The chemical potential for a macroscopic salt concentration can be reliably determined

The salt insertion free energies and average volumes in Figure 4 *upper left* provide a way to relate the chemical potential Δ*µ* to macroscopic salt concentration ⟨*ĉ*⟩ via equation 19. Figure 4 *upper right* shows the predicted macroscopic salt concentration for a range of chemical potentials Δ*µ* computed using equation 19. The average salt concentration in a saline box ⟨*c*⟩ should equal the predicted macroscopic concentration for sufficiently large saline boxes if the chemical potential has been properly calibrated. To verify the accuracy of the calculated values for *f*(*N*_NaCl_) and 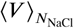, simulations of water boxes, that initially had no salt present, were performed using an osmostat with different fixed chemical potentials and the average salt concentrations in the simulations were estimated (Figure 4; *upper right*). These boxes of TIP3P and TIP4P-Ew waters contained 4085 and 4066 molecules respectively, whereas the TIP3P and TIP4P-Ew boxes used to estimate *f* (*N*) and 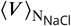 contained 887 and 886 molecules respectively. As Figure 4 *upper right* shows, the macroscopic concentrations ⟨*ĉ*⟩ predicted using equation 19 fall within the statistical error of the average concentrations ⟨*c*⟩ determined from the fixed-Δ*µ* simulations.

Although Δ*µ* is the thermodynamic control parameter for osmostated simulations, experimental wetlab conditions instead generally specify the macroscopic salt concentration ⟨*ĉ*⟩ rather than Δ*µ*. As the relationship between Δ*µ* and ⟨*ĉ*⟩ is monotonic, as illustrated by Figure 4 *upper right*, we can numerically invert equation 19 to enable practitioners to choose the desired macroscopic salt concentration and extract the required Δ*µ* for the osmostat to model equilibrium with the macroscopic salt concentration ⟨*ĉ*⟩.

### The average salt concentration is highly sensitive to chemical potential

The macroscopic salt concentration ⟨*ĉ*⟩_Δ*µ*_ for a fixed chemical potential Δ*µ* is a highly sensitive and non-linear function of the chemical potential (Figure 4; *upper right*) for both water models. Small changes to the chemical potential, on the order of 1 *kT*, can alter the mean concentration by hundreds of millimolar. Correspondingly, to accurately model a given macroscopic concentration *c*, the function Δ*µ*(*c*) must be very precisely calibrated.

### Different water models have distinct chemical potentials for the same salt concentration

Strikingly, both the value and shape of ⟨*ĉ*⟩_Δ*µ*_ is very sensitive to choice of water model (Figure 4; *upper right*). For instance, a Δ*µ* of about 316 *kT* results in a mean salt concentration in TIP3P water that is approximately 500 mM, compared to approximately 200 mM in TIP4P-Ew water for the same value of Δ*µ*. These features highlight the importance of specifically calibrating the chemical potential for each water and ion model as well as estimating *f*(*N*_NaCl_) and 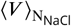 to a sufficient degree of precision. Figure A5.2 shows that for TIP3P and the treatment of long-rang interactions used herein, the free energies *f*(*N*_NaCl_) for each *N*_NaCl_ ∈ {0, 1, …, 20} need to be determined to a standard error of 4 kcal/mol to consistently determine the macroscopic concentration to an inaccuracy of at least about 80 mM for 1 mM ≤ ⟨*ĉ*⟩ ≤ 1000 mM. The average standard error achieved in the calibration simulations for the free energies *f*(*N*_NaCl_) is 0.02 kcal/mol, which determines the concentration to an inaccuracy no larger than about 1 mM.

### NCMC greatly enhances the sampling efficiency of salt insertion and deletion moves

We estimate that instantaneous salt insertion and deletion moves have acceptance probabilities of 3.0× 10^−51^ [95% CI: 5.0 × 10^−66^, 9.0 × 10^−51^] and 1.0 × 10^−46^ [95 % CI: 3.0 × 10^−64^, 4.0 × 10^−46^] in TIP3P and TIP4P-Ew water respectively, implying that the implementation of an osmostat is practically impossible using such naïve moves. In contrast, we found that in our longest protocol, NCMC insertion/deletion attempts achieved acceptance probabilities of about 30% in TIP3P water and approximately 15% in TIP4P-Ew water (see the lower left of Figure 4). Although the acceptance probability increases monotonically with the length of the protocol, so does the computational cost and time for each attempt. The efficiency, defined in equation 25, quantifies the trade-off between the acceptance rate and computational expense. Figure 4 *lower right* shows that NCMC protocols in TIP3P water that are between 15 ps and 30 ps in length are the most efficient for our procedure. For this reason, all subsequent simulations used TIP3P water and a 20 ps long NCMC protocol. In addition, it was found that 10 propagation steps (at 2 fs) between each perturbation was found to be the most computationally efficient for our simulation code SaltSwap [0.5.2] and OpenMM [7.1.0] (see Figure A5.3). Further optimization of the NCMC protocol would be required for NCMC attempts in TIP4P-Ew to achieve sampling efficiencies that are competitive with those in TIP3P water.

### An NCMC osmostat can rapidly equilibrate the salt concentration in biomolecular systems

Figure 5 shows example salt concentration trajectories around DHFR as well as plots of the corresponding autocorrelation functions for three biologically plausible macroscopic salt concentrations. The autocorrelation times for the three macroscopic salt concentrations are on the order of 1 ns, implying that our osmostated simulations should be at least tens of nanoseconds long to generate sufficient uncorrelated samples of salt concentrations. Importantly, the magnitude of the instantaneous salt concentration fluctuations increases with the macroscopic salt concentration, which causes an increase in the correlation time as our osmostat implementation proposes the insertion/deletion of one salt pair a at a time. As a result, more attempts are required to explore salt concentration distributions of higher variance. This suggests that inserting or deleting multiple salt pairs in each attempt could improve the sampling efficiency of our osmostat at higher macroscopic salt concentrations, though longer NCMC insertion/deletion protocols would likely be required to achieve similar acceptance probabilities.

**Figure 5.**
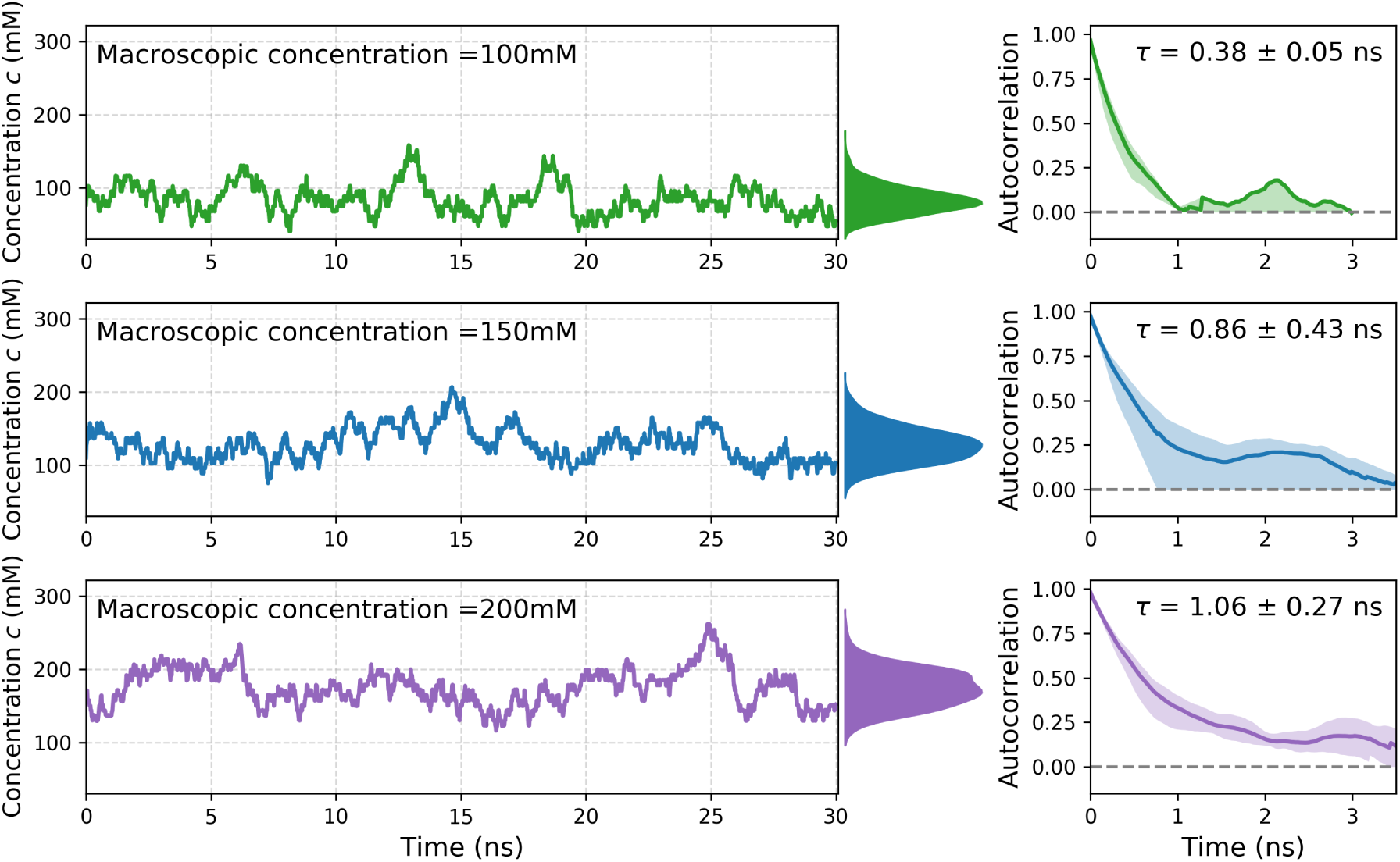
Dynamic salt sampling for DHFR in TIP3P water at three macroscopic salt concentrations. *Left:* Trajectories of the salt concentration in 30 ns simulations of DHFR in a boxes of TIP3P waters as a function of time for 100 mM, 150 mM, and 200 mM NaCl, along with distribution of equilibrium salt concentrations to right of the time-series plots. The distributions were estimated using a Gaussian smoothing kernel with bandwidth of 0.3 mM from all three simulation repeats at each macroscopic concentration. Before the insertion of NaCl, the simulation contained 7023 water molecules. *Right:* Normalized fluctuation autocorrelation functions and integrated autocorrelation times (*τ*) of salt concentrations for each simulation. Shaded regions and uncertainties on the autocorrelation time signify 95% confidence intervals calculated using bootstrap estimation from three independent simulations.

### Fluctuation magnitude grows with system size and macroscopic salt concentration

Figure 6 *upper left* demonstrates that for a pure box of saline and fixed macroscopic salt concentration, increasing the number of molecules in the system increases both amount of salt and the spread of the salt number distribution; in contrast, Figure 6 (*upper right*) reveals that the distribution of the concentration remains centered around the macroscopic concentration, but the variance decreases. Both of these trends are to be expected from statistical mechanics (see Appendix 2). The salt concentration distribution for the smallest water box (with 2094 molecules) in Figure 6 (*upper right*) can be seen to be highly multimodal. Each peak corresponds a particular number of salt pairs in the system; there are so few water molecules in this system that changing *N*_NaCl_ by one results in a large jump in the concentration. Figure 6 (*bottom left and right*) highlight that for a system with a fixed number of water molecules, the number of salt pairs increases in proportion with the macroscopic concentration.

**Figure 6.**
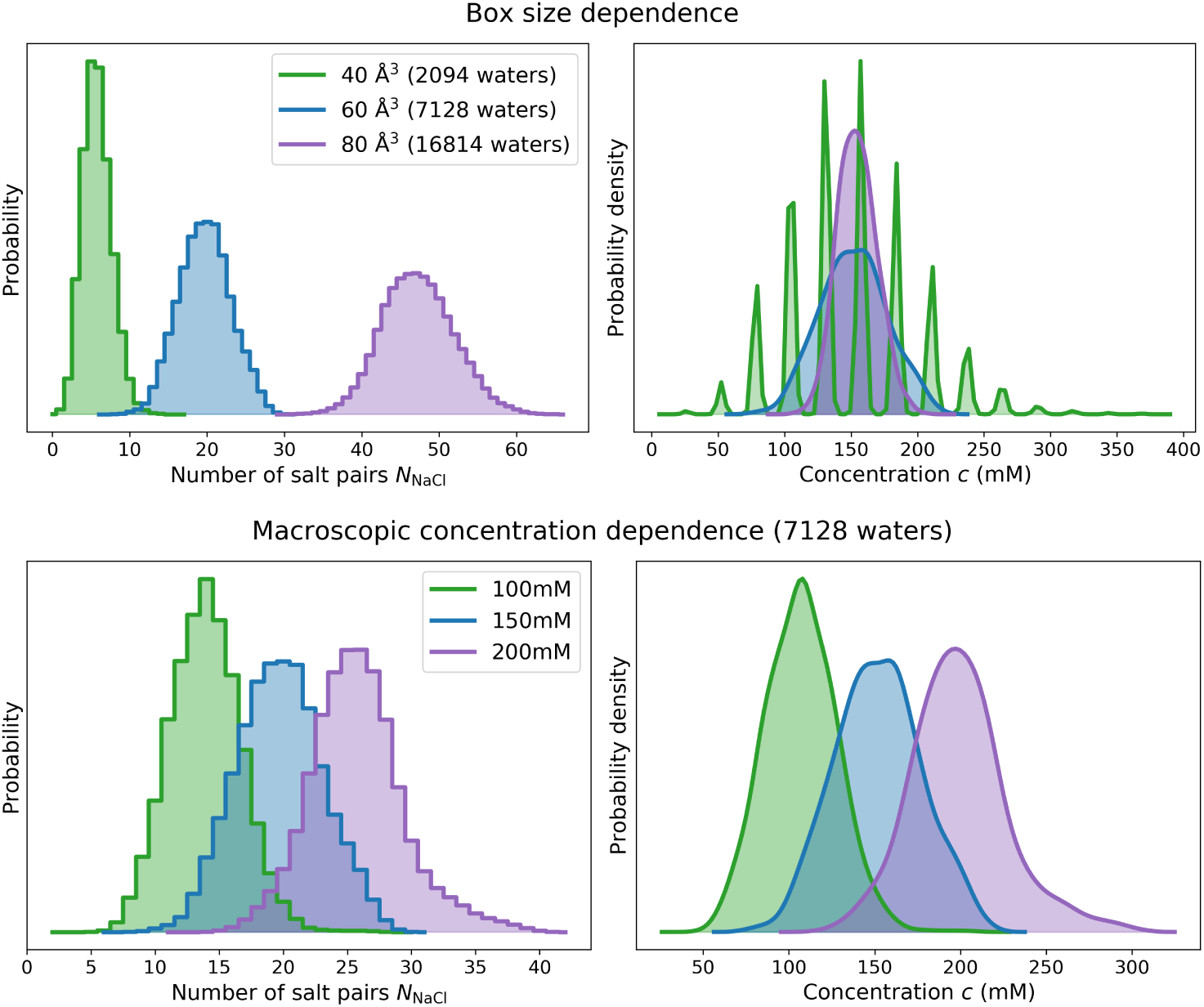
Distribution of salt numbers and concentrations for TIP3P water boxes of varying size and macroscopic salt concentration *Top:* Equilibrium distribution of salt numbers (*N*_NaCl_, *left*) and salt concentrations (*c, right*) as a function of the number of water molecules in the simulation. The applied macroscopic concentration was 150mM. blueAs expected (see Appendix 2), at fixed macroscopic salt concentration, the magnitude of fluctuations in the number of salt pairs *N*_NaCl_ grows with box size (*left*), whereas the magnitude in the concentration decreases with box size. The average salt concentration ⟨*c*⟩ remains fixed at the specified macroscopic concentration (*right*) showing that the calibrated chemical potential Δ*µ* is invariant to box size provided the calibration box is selected to be sufficiently large to avoid finite-size effects. The small range of *N*_NaCl_ in the 40 Å box results in a multimodal salt concentration distribution. *Bottom:* Equilibrium distribution of salt numbers (*N*_NaCl_, *left*) and salt concentrations (*c, right*) as a function of salt concentration for a water box containing 7128 waters.

### Salt concentrations vary significantly in typical biomolecular systems

Figure 7 shows the distribution of salt concentration and ionic strength for 3 typical biomolecular systems: DHFR, *apo* Src kinase, and the Drew-Dickerson DNA dodecamer. The distributions in a box of TIP3P are also shown for reference. The fluctuations of the salt concentration around the macromolecules are substantial: 95% of all salt concentration samples fall within a range of 90.2 mM for DHFR, 87.7 mM for Src kinase, and 135.6 mM for the DNA dodecamer system. We expect these values to be indicative of the natural variation in salt concentration in the local environments of real biomolecules.

**Figure 7.**
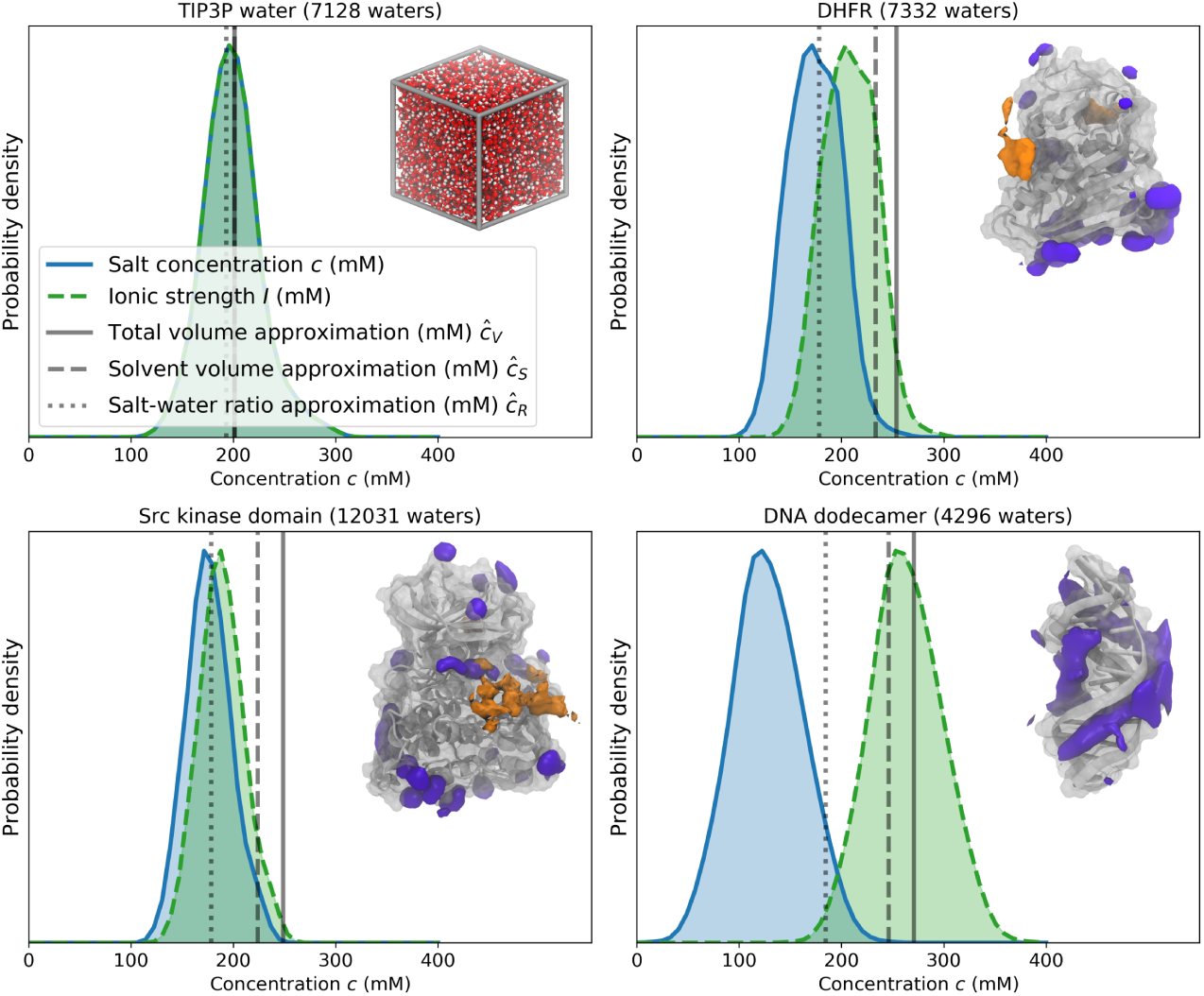
Equilibrium salt concentration distributions for various biomolecular systems simulated with a 200 mM osmostat. Equilibrium salt concentration distributions (blue shaded area) are shown as a kernel density estimate of the probability density, along with the ionic strength of the solvent (light green shaded area with dotted lines). No samples of the salt concentration were discarded for these density estimates. For reference, the mean salt concentrations that would be achieved in three typical fixed-salt salination strategies are shown in transparent gray lines. The continuous line uses equation 22 and the *total* volume of first frame of the production simulation; the dashed line uses equation 23 and the volume of *solvent* at the start of the production simulation, and the dotted line uses equation 24 and the *ratio* of the number of salt pairs and water molecules. Illustrations of each system are also shown in the top right of each plot, with Na+ (purple) and Cl- (orange) densities from equilibrium 200 mM osmostat simulations shown around the three macromolecules. Isovalues for the each of 3D ion densities were chosen for visual clarity. *Upper left:* Box of TIP3P waters; *Upper right:* DHFR (dihydrofolate reductase) in TIP3P with isosurfaces containing 14.3% and 0.8% of Na^+^ and Cl^−^ densities, respectively; *Lower left: apo* Src kinase in TIP3P with isosurfaces containing 8.5% and 0.6% of the Na+ and Cl-densities, respectively; *Lower right:* Drew-Dickerson DNA dodecamer in TIP3P with 8.9% of the Na^+^ density contained in the isosurface.

### Simulations containing charged biomolecules can experience salt concentrations that deviate systematically from the macroscopic concentrations

The DHFR, *apo* Src kinase, and the Drew-Dickerson DNA dodecamer structures have net charges of -11 |*e*|,-6 |*e*| and -22 |*e*|, respectively. The net charge of the DNA dodecamer is a result of the phosphate group on each of the nucleotides (with each of the eleven phosphate groups carrying -1 |*e*| charge), whereas the net charges on DHFR and Src kinase are due to an excess of glutamate and aspartate residues over arginine, histidine, and lysine residues. Neutralizing Na^+^ ions were added to both systems to avoid the uniform background charge that would be applied automatically with PME electrostatics. Like the other ions in our osmostat, these counterions had transmutable identities.

Figure 7 shows that in our osmostated simulations of the macromolecules, the average salt concentration is on average *less* than the macroscopic salt concentration. This is particularly apparent with the DNA dodecamer, which has a mean concentration of 128.0 [121.5, 134.5] mM (where the quantity in brackets denotes the 95% confidence interval of the mean concentration). The salt concentration distribution in the DHFR and Src kinase systems are centered closer to the macroscopic concentration of 200 mM, with estimated means of 174.0 [164.4, 180.4] and 176.3 [171.6, 189.5] mM, respectively. To compute these statistical estimates and confidence intervals, no data was discarded at the start of the simulation, and approximately statistically independent concentration samples were extracted using the pymbar timeseries module^75^.

The larger number of water molecules in the Src kinase system is partly the reason why its mean concentration is closer to the macroscopic value than the DNA dodecamer. Bulk-like conditions anchor the sampled salt concentrations about the macroscopic concentration; the more water molecules and salt pairs there are, the smaller the effect a macromolecule has on the salt concentration relative to the whole system. Figure 8 *inset* highlights this phenomenon with the DNA dodecamer; the mean salt concentration moves closer to the macroscopic value when more water molecules are added to the simulation.

**Figure 8.**
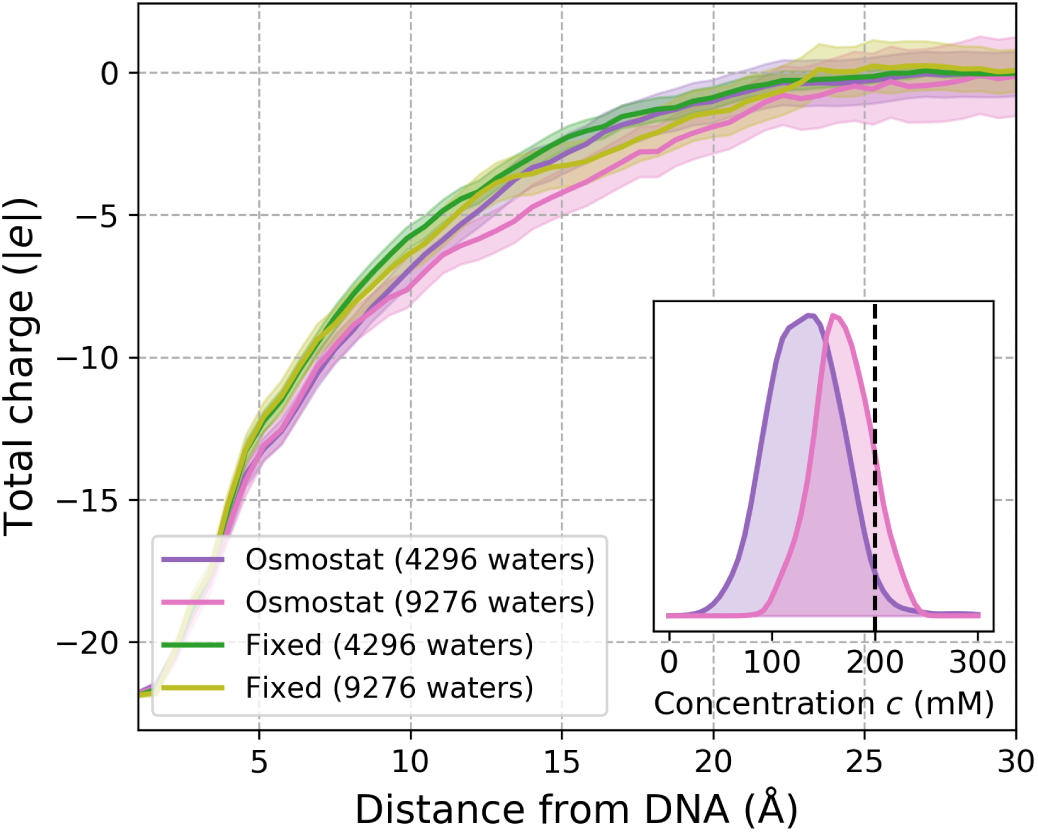
Dependence of the charge screening length and salt concentration on simulation size for the Drew-Dickerson DNA dodecamer. *Main:* The mean total charge within a minimum distance from the Drew-Dickerson DNA dodecamer for 200 mM NaCl osmostated simulations and 200 mM fixed salt fraction simulations. To compare the effect of solvent content on charge screening effects, the DNA dodecamer was solvated in water boxes of two different sizes. The smallest system had water added up to a distance no less than 9 Å away from the DNA dodecamer (adding 4296 waters), whereas the larger was solvated up to a distance at least as large as 16 Å (adding 9276 waters). As each simulation is electrostatically neutral, the total charge must decay to zero as the distance from the DNA dodecamer increases, but the rate at which this decay occurs provides insight into the lengthscales for which biomolecules accrete a neutralizing ion constellation. The charge distributions appear robust with respect to the size of the simulation cell, as all 95% confidence intervals (transparent colors) of the mean charge-distance profiles overlap over all distances considered. The charge-distance profiles were estimated by counting the number of ions within fixed distances of the DNA dodecamer every 1 ns and the confidence intervals were estimated by using boostrap sampling. *Inset:* Salt concentration probability densities estimated using kernel density estimation for 200 mM osmostated simulations with different amounts of solvent. The simulation with the small solvent box (purple) recruits far fewer salt pairs from bulk on average (dotted black line denotes 200 mM), while the average salt concentration of the simulation with the larger solvent box (pink) is significantly less perturbed from bulk.

### The accuracy of heuristic salination schemes is system dependent

On its own, the excluded volume of the macromolecule will reduce the number of salt pairs that can occupy the simulation volume compared to bulk saline. So, as we define the salt concentration as the number of salt pairs over the total volume of the system (equation 20), one would expect there to be a lower salt concentration than the macroscopic value. The preparation schemes that are typically used to add salt in fixed-salt simulations that account for this effect use either the volume of the solvent (equation 23), or the ratio of the number of salt pairs to water molecules (equation 24). As a result, these methods are closer to the mode of the concentration distributions in the osmostated simulations than the heuristic method that uses the total volume of the system (equation 22). The volume-based methods are sensitive to how equilibrated the volume is when salt is added, and, in Figure 7, the volume at the start of the production simulation was used to estimate the amount of salt that would be added with equations 22 and 23. The salt-water ratio method (equation 24) has no such volume dependence, which is partly why it is a better predictor for the salt concentration than the others.

### The ionic strength exceeds the salt concentration for charged macromolecules

In addition to the distributions of salt concentrations, Figure 7 also shows the ionic strength of the saline buffer. While the ionic strength is used in analytical models to estimate the activities of ionic species^83^, the only discernible common feature of the ionic strength in our simulations is that it tends to be greater than the salt concentration, which is predominantly due to the presence of neutralizing counterions. The estimated mean ionic strength of the saline buffer in the macromolecular systems are 208.2 [198.2, 213.6] mM for DHFR, 189.0 [179.5, 196.4] mM for Src kinase, and 263.4 [256.6, 269.8] for the DNA dodecamer. It is important to note that the calculated ionic strength can be much larger when the contribution of the macromolecule is included: the estimated ionic strengths for the whole of the DHFR, Src kinase, and DNA systems are 551.0 [541.0, 556.4] mM, 263.6 [253.8, 270.8] mM, and 3241.6 [3227.3, 3244.7] mM respectively. These high values, particularly for the DNA system, is because the ionic strength is proportional to the square of the charged number of the ionic solute. It could be more informative to consider the macromolecule and the counterions that are bound to it as a single, aggregate macro-ion, such that the contribution to the ionic strength would be lessened^83^; however, as there is no clear boundary between bound and unbound ions (see Figure 8), this approach is conceptually difficult.

### The osmostat accurately represents the local salt concentration around DNA

The aim of our osmostat is to replicate the local ion concentrations that would occur around biomolecules when embedded in large saline reservoirs. However, the use of periodic simulation cells and the addition of neutralizing counterions constrains length scale at which charges are screened (the Debye length) to be less than or equal to the length scale of the periodic cell. An artificial constriction of the Debye length would be finite size effect that would limit the accuracy of the salt concentrations from osmostated simulations. Figure 8 shows the total charge contained within ever increasing distances from the Drew-Dickerson DNA dodecamer for two simulation box sizes. The smallest box was constructed by solvating the DNA up to a minimum distance of 9 Å away from the DNA (4296 water molecules), whereas the larger box resulted from solvating up to a distance of 16 Å from the DNA (9276 water molecules). If the Debye length was significantly affected by the periodic cell size of the smallest simulation, there would be large discrepancies between the charge distributions around the DNA of the smallest box and the larger box. Figure 8 indicates that if such discrepancies exists, they are small, and are not found to be statistically significant in our analysis.

Shown first in Figure 7 (lower right), the osmostated simulation of the Drew-Dickerson DNA dodecamer experienced significantly lower NaCl concentrations than the applied 200 mM macroscopic NaCl concentration. This difference highlights how the local ionic environment of a solute can be strikingly different from bulk saline. Increasing the amount of water in the simulation diminishes the relative effect that DNA has on perturbing the salt concentration distribution of the whole system. Figure 8 (inset), shows that increasing the number of water molecules in the system from 4296 to 9276 molecules partially masks the local salt concentration around the DNA, such that the total salt concentration over the whole system is closer to the macroscopic concentration of 200 mM.

### The NCMC osmostat can efficiency of ion-biomolecule interactions

To compare the computational efficiency of NCMC ion sampling to that of fixed-salt MD simulations, the autocorrelation functions of cation-phosphate interactions were estimated from the DNA dodecamer simulations. Cation-phosphate interactions were recorded as every time a cation was within 5 Å of the phosphorous atoms in adenine nucleotides. This cutoff was chosen following the DNA convergence analysis of Ponomarev et al.^25^. The autocorrelation function of these interactions measures the probability that a cation that is initially within the distance cutoff will also be present after a given amount of time. As our osmostat uses NCMC to add and remove ions, one would expect the osmostat interaction autocorrelation function to decay significantly faster than that from the fixed salt simulations when only considering the molecular dynamics—Figure 9 shows that this is indeed the case.

**Figure 9.**
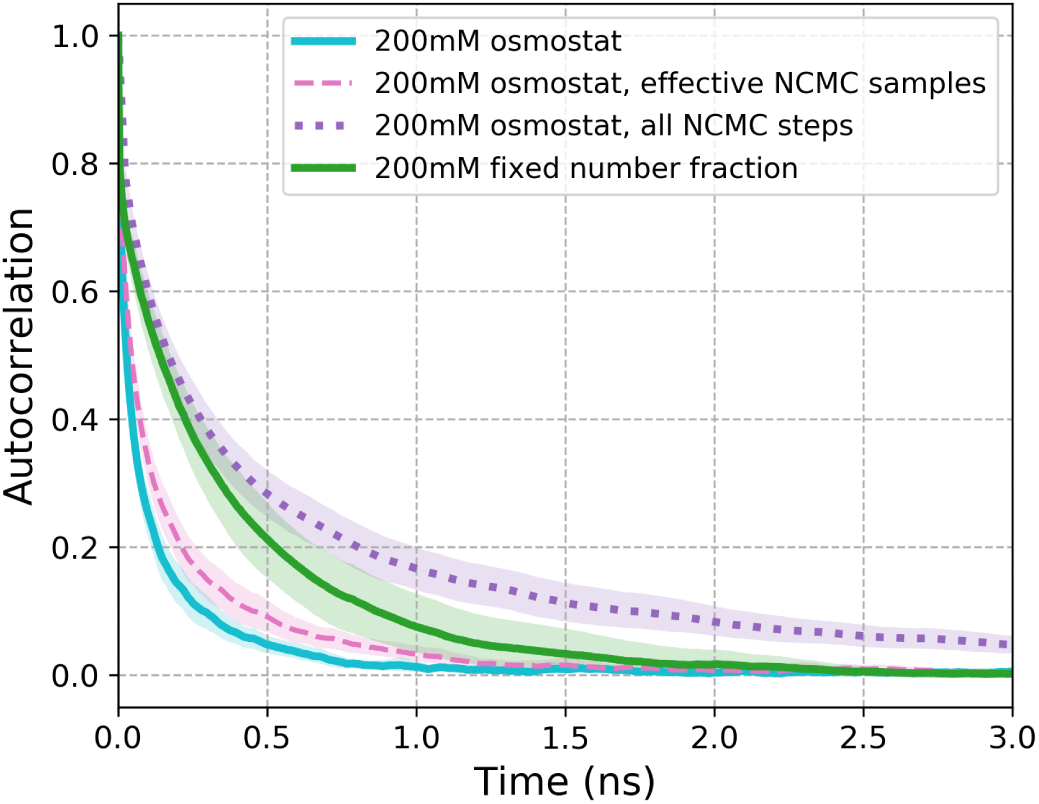
Phosphate-cation normalized fluctuation autocorrelation functions for binary occupancies around a DNA palindrome. The Drew-Dickerson DNA dodecamer (CGCGAATTGCGC) is a palindromic DNA sequence that has been traditionally been used as a demonstration of the slow convergence of ion distributions around the phosphate backbone of DNA. Phosphate-cation normalized fluctuation autocorrelation functions for binary occupancies in standard MD (thick green) and MD with dynamic ion sampling either neglecting the NCMC switching time (thick cyan), or the effective number of samples taken with accepted NCMC moves (dashed pink), or accounting for all NCMC MD steps whether the moves were accepted or not (dotted purple). The latter accounts for the total computational expense of our NCMC protocol. Shaded regions highlight 95% bootstrap confidence intervals, with bootstrap samples taken from all the adenine groups from the three simulations.

When the simulation time from NCMC is not considered, the phosphate-cation interaction autocorrelation function from the osmostat simulations decays significantly faster than the fixed salt simulations (Figure 9). The corresponding integrated autocorrelation times for osmostated simulations and fixed-salt simulations are 0.11 [0.09, 0.13] ns and 0.29 [0.23, 0.36] ns respectively. As each accepted NCMC move has propagated the configurations of the whole system, the faster decorrelation of DNA-ion interactions could be a result of these extra propagation steps, as opposed to the fact that ions are being inserted and deleted. As described in the methods, a salt insertion or deletion attempt occurs every 4 ps, and an NCMC attempt involves 20 ps of dynamics. The average acceptance probability in the DNA simulations was calculated to be 11.9 [11.7, 12.2] %. Therefore, the osmostated simulations propagate the system 1.6 [≈(0.119× 20 ps + 4 ps)/4 ps] times as much dynamics than fixed salt simulations. Multiplying the osmostated integrated autocorrelation time by this factor results in a value that remains significantly less than the integrated autocorrelation time from the fixed salt simulations. Figure 9 *right* shows the osmostated autocorrelation function when the timescale has been multiplied by the effective NCMC sampling factor (1.6). Despite the application of this factor, the fixed-salt autocorrelation function can be seen to decay significantly slower than the stretched osmostated autocorrelation function. Thus, the increased sampling efficiency observed in the osmostated simulations cannot be explained by the extra dynamics sampled in the NCMC simulations. This implies that the random insertion and deletion, not the NCMC that was used to enhance the move efficiency, is responsible for the rapid decorrelation of ion interactions observed in the DNA osmostated simulations.

The total number of NCMC timesteps (including from rejected moves) can be used to account for the additional computational burden of the NCMC osmostat in the phosphate-cation autocorrelation times. There is an additional 20 ps of dynamics for every insertion/deletion attempt, irrespective of whether the proposal was accepted or not. As each attempted is preceded by 4 ps of equilibrium dynamics, our osmostated simulations have 6 (= (20 ps + 4 ps)/4 ps) times as timestep evaluations than the fixed-salt simulations. Multiplying the mean integrated autocorrelation time from the osmostat simulations by this factor yields an effective autocorrelation of 0.65 [0.55, 0.75] ns. Although this estimate now exceeds the upper confidence interval of the fixed-salt integrated autocorrelation time (0.29 [0.23, 0.36] ns), there is only approximately 0.1 ns difference between the lower and upper confidence intervals. Figure 9 also shows the osmostat phosphate-ion autocorrelation function when the all the NCMC propagation steps (including rejected moves) are accounted for. One can see that for below ∼1 ns, the 95% confidence intervals of the autocorrelation functions overlap with those of fixed-salt autocorrelation function. These results imply the dynamic NaCl sampling achieved by our osmostat has a similar cost effectiveness—with regards to ion sampling—than fixed-salt simulations, with the additional benefit of sampling realistic salt concentrations.

## Discussion

In this work, we have implemented an osmostat that dynamically samples the NaCl concentration in biomolecular simulations. The osmostat couples a simulation cell to a saline reservoir at a fixed macroscopic concentration and allows the salt concentration in the simulation to fluctuate about its equilibrium value. We have applied our osmostat to simulations of dihydrofolate reductase (DHFR), *apo* Src kinase, and the Drew-Dickerson B-DNA dodecamer (CGCGAATTGCGC), and found that the mean salt concentration can differ significantly from the amount salt added by common molecular dynamics methodologies. In addition, we found that the salt concentration fluctuations were large, being of the same order of magnitude as the mean. These results show that the ionic composition around biomolecules can be highly variable and system dependent.

The insertion and deletion of salt was greatly enhanced by nonequilibrium candidate Monte Carlo (NCMC), to the extent that the protocol used in our simulations was approximately 5 × 10^46^ times more efficient than instantaneous attempts in TIP3P water. The Drew-Dickerson B-DNA dodecamer is a palindromic sequence that facilitated a study of the convergence of ion distributions around the DNA. We found that, despite the additional computational expense of the NCMC osmostat, the sampling and computational efficiency of DNA:ion interactions remained comparable to fixed-salt simulations. However, it is important to note that made no effort to optimize the NCMC protocols beyond selecting an appropriate total switching time for NCMC moves—it is possible that further optimization of these protocols using recent techniques based on mapping geodesics in the thermodynamic metric tensor space^84–88^ can lead to increased efficiency.

### Potential applications

While the dependence of enzyme-substrate activity on ionic strength is well documented, the impact of salt concentration on protein-ligand binding affinity is much less clear. Recently, Papaneophytou et al. performed a systematic analysis on the effect of buffer conditions on the *in vitro* affinity of three complexes^15^, finding salt concentration dependence to be system dependent and largest for complexes that formed hydrophilic interactions. Our osmostat provides the opportunity to rigorously study the impact of salt concentration on protein-ligand binding affinities *in silico*. We are interested to know if similar trends to what Papaneophytou et al. observed can be reproduced in all-atom binding free energies calculations, and whether binding free energy estimates differ significantly between simulations carried out with and without an osmostat. Free energy calculations on complexes whose association is sensitive to the concentration of salt are likely to be most affected by the osmostat, given the large fluctuations of concentration and the deviation from the fixed-salt values that occurred in our simulations (see Figure 7). The combination of self adjusted mixture sample (SAMS) and Bennett acceptance ratio (BAR) that we used to calibrate the chemical potential can also be used to estimate the difference between traditional and osmostated free energy calculations. If significant differences between binding free energy calculations in fixed-salt and osmostat simulations are observed, it is also possible to apply the same SAMS-BAR methodology to correct the free energy calculations that have been performed with fixed salt.

As our osmostat has been designed to reproduce realistic salt environments around biomolecules, it is well suited to study systems whose function are sensitive to the salt concentration, or biomolecules that are regulated by interactions with Na^+^ or Cl^−^. While our osmostat can efficiently sample ion binding to biomolecular surfaces, the sampling of deeply buried ion binding sites is likely to be no more than efficient than in typical molecular dynamics simulations due to the fact that our osmostat is implemented by swapping water with salt. To this end, the osmostat could be improved and generalized if position-biased insertions of fully-decoupled ghost molecules could be added to its sampling repertoire. An example of one such biasing scheme can be found in the biomolecular simulation package ProtoMS, where the grand canonical insertion and deletion of water are attempted in a pre-defined region within proteins^48,49^. Previously, Song and Gunner studied the interplay between protein conformation, residue pKas, and ion binding affinity using a grand canonical ion insertion scheme within the MCCE framework^6^. Their work provided structural insight into the often tight-coupling between ion and proton affinity as well as the pH sensitivity of ion binding, and highlights the power of specialized ion sampling schemes to rationalize and understand experimental measurements. The insertion of decoupled ghost molecules—while it would likely require more highly optimized alchemical protocols for insertion—would also permit generalizing the method to more complex salt or buffer molecules or other excipients.

### Enhancing realism in molecular simulations

Because the pKa of protein residues are dependent on the ionic strength of the medium, a natural extension of the osmostat is to combine it with constant-pH simulations in explicit water. Previously, Chen and Roux coupled protonation state changes with the insertion and deletion of ions to maintain electrostatic neutrality^33,34^. The application of an osmostat to such transformations would allow for the macroscopic ion concentration—as well pH—to be rigorously maintained, and could be implemented in modular MCMC scheme that updates protonation states and ion identities in tandem.

This work only considers the concentration of NaCl, but both the formalism we introduce in the Theory section and the flexibility SaltSwap code-base can be readily extended to sample over biologically relevant salt mixtures by including additional monovalent species such as K^+^ and divalent species like Ca^2+^. More complex ions or buffer molecules, such as 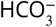 would require a more significant extension to code (such as the insertion of ghost particles described earlier), and could be implemented by using a softcore alchemical NCMC pathway that converts the molecule between fully interacting and noninteracting states.

The combination of a multicomponent osmostat with a constant-pH methodology would allow for realistic physiological conditions to be better approximated in molecular simulations. While it is well appreciated that pathological tissue can be found with altered pH—tumor microenvironments can have low pH, while cancer cells can have elevated pH, for example^89^—pathologies can also disrupt healthy ion compositions^5^. The ability to reproduce specific ionic concentrations as well as pH would open the possibility of using molecular simulations to target compounds to specific microenvironments or achieve selectivity via salt-dependent environmental differences. Indeed, Spahn et al. recently used molecular simulations to develop an analgesic that selectively targets the *µ*-opioid receptors in damaged, low pH, tissues^90^.

### Improving osmostat efficiency

We have demonstrated that our implementation of the NCMC osmostat was sufficient to sample equilibrium distributions of ions around biomolecules in practical simulation times. We have not yet extensively optimized the osmostat for computational or algorithmic efficiency beyond exploring NCMC protocol lengths (Figure 4 and Figure A5.3), such that there a number of ways that the computational efficiency could be further improved.

In our current implementation, which only proposes insertion/deletion of a single salt pair in each proposal, the correlation time for the instantaneous salt concentration increases with increasing system size as the size of the equilibrium fluctuations also grow in terms of total numbers of ions (Figure 5). Inserting or deleting multiple ion pairs—likely using longer specialized NCMC protocols tuned to the number of ions being inserted or deleted—could help maintain efficiency. Adaptive MCMC proposals, currently in widespread use in the Bayesian inference community (e.g., PyMC^91^), could be used to automatically tune the number of ions proposed to be deleted or inserted based on the current concentration and the history of the sampler, provided care was taken to ensure the adaptation method maintained ergodicity and ensured the target density was properly sampled^92^. One of the earliest adaptive scheme was originally validated on unimodal distributions^93^, such that a discretized variant could be well suited to sampling the number of pairs.

Acceptance rates can also be increased by using proposals that do not simply select ions at random, but instead select ions that are more easily inserted/deleted based on some rapidly-evaluated surrogate (such as their instantaneous Monte Carlo acceptance probabilities or the electrostatic potential on water and ion sites), provided this biased selection probability is accounted for in a modified Metropolis-Hastings acceptance criteria.

There is a great deal of potential to improve the efficiency of the NCMC protocol used for the insertion and deletion proposals. The current work uses a linear interpolation of the salt and water nonbonded parameters as the alchemical path and perturbations steps that are equally spaced with respect to the parameters, primarily because this is the simplest scheme to implement. The only optimization carried out here was tuning the total protocol length to be sufficiently long to achieve high acceptance rates but not so long that the overall efficiency would be diminished by further extending the protocol length (Figure 4). Optimized NCMC protocols can reduce protocol switching times required to achieve high acceptance rates, thereby increasing overall efficiency. The ability to quantify the *thermodynamic length* of the nonequilibrium protocol allows the problem of protocol optimization to be tackled rigorously. The thermodynamic length (an application of the Fisher-Rao metric to statistical mechanics^94^) is a natural, albeit abstract, measure of the distance traversed by a system during a thermodynamic driving process^84^.

Within this framework, optimal NCMC protocols are given by geodesics in a Riemannian metric tensor space^86^. The thermodynamic length of the NCMC protocol can be estimated in separate equilibrium simulations spaced along the alchemical path, or estimated directly from the protocol work values of the NCMC switching trajectories, including those from rejected proposals^85^. For optimizing a preselected alchemical path, spacing the perturbation steps to be equidistant with respect to the thermodynamic length can improve acceptance rates by reducing the total variance of the protocol work. As optimal paths are geodesics in thermodynamic space, the most efficient alchemical path for the insertion or deletion will likely be a nonlinear, rather than linear, interpolation of the water and ion nonbonded parameters. Previous efforts to optimize nonequilibrium paths have included directly solving for the geodesic^87^, sampling the protocol from an ensemble^88^, and by restricting the optimization to a family of functional forms^95^. The close relationship between thermodynamic length and the dissipation along the path also suggests that restricting the propagated dynamics to only the first few layers of the solvation shell around the transmuted molecules could also improve the NCMC protocol.

### Conclusion

The philosophy of this work is that increasing the realism of biomolecular simulations will aid structural inference and improve the quantitative accuracy of predictions. We believe that the NCMC osmostat we have presented here will be a useful tool for probing the interactions of ions and biomolecules under more physiological conditions than considered in traditional molecular dynamics simulations. It is our hope that the application of the osmostat to protein-ligand binding free energy calculations and extending the method to more comprehensive ion compositions will improve its utility even further.

## Code and data availability

- Code is available at https://github.com/choderalab/saltswap
- Data analysis scripts available at https://github.com/choderalab/saltswap-results

## Acknowledgments

GR, ASR, PBG, JF, and JDC acknowledge support from the Sloan Kettering Institute. JDC acknowledges support from NIH grant P30 CA008748. JF acknowledges support from NSF grant CHE 1738979. PBG acknowledges support from Silicon Therapeutics Open Science Fellowship. The authors are especially grateful to Zhiqiang Tan (Rutgers) for many fruitful discussions on the theory and application of self-adjusted mixture sampling and nonequilibrium candidate Monte Carlo, to Marilyn R. Gunner (CCNY) for her insight in counterion distributions around macromolecules, Peter Eastman (ORCID: 0000-0002-9566-9684) for his help and advice with OpenMM, and Kyle A. Beauchamp (ORCID: 0000-0001-6095-8788) for the DHFR system. JDC expresses enormous gratitude to Ken A. Dill (ORCID: 0000-0002-2390-2002) for highly formative conversations about the nature of thermodynamic parameters relevant to biomolecular systems on microscopic lengthscales and his encyclopedic knowledge of the foundational experiments that laid bare their role in driving biophysical phenomena; some of these ideas are realized here.

## Appendix 1

### Symbols and their definitions

- *x* : Instantaneous configuration (positions, box vectors)
- 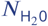: Number of water molecules
- 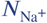: Number of cations
- 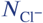: Number of anions
- *N*_NaCl_ : Number of salt pairs beyond minimal neutralizing ions; equal to min 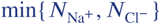
- *N* : Sum of total number of waters and ions in the system
- *θ* : Vector species labels with *N* elements that identifies which molecules are waters and which are ions; *θ*_*i*_ = 0 indicates water, *θ*_*i*_ = +1 indicates monovalent cations, and *θ*_*i*_ = −1 indicates monovalent anions
- *z* : total charge number of the macromolecules in the simulation
- *n*(*θ*) : total charge number of the ions in the simulation

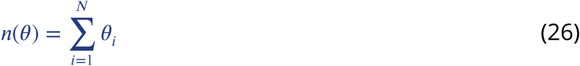
- *U* (*x, θ*) : Potential energy for a system with configuration *x* and water/ion identities *θ*, units of energy
- *p* : External pressure, units of energy · length^−3^
- *V* : Instantaneous box volume, units of length^3^
- *T* : Absolute temperature, units of temperature
- *k*_*B*_ : Boltzmann constant, units of energy · temperature^−1^
- *β* : Inverse temperature (≡ 1/*k*_*B*_*T*), units of energy^−1^
- *I* : Ionic strength, where instantaneous ionic strength for configuration *x* is given by

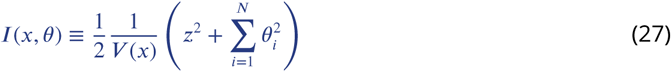 Note that ionic strength includes minimal neutralizing counterions in the sum.
- Δ*µ* : Chemical potential difference for extracting a NaCl molecule from bulk water and depositing two water molecules to bulk water; an abbreviation of 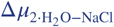
- *f*(*N*_NaCl_) : Free energy to replace 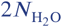 water molecules with *N*_NaCl_ salt pairs in bulk water; an abbreviation of *f*(*N*_NaCl_, *N, p, T*).
- Δ*f*(*N*_NaCl_) : Free energy to add one more salt pair and remove two additional water molecules in a box of water than contains *N*_NaCl_ salt pairs already; equal to *f*(*N*_NaCl_ + 1) − *f*(*N*_NaCl_); an abbreviation of Δ*f*(*N*_NaCl_, *N, p, T*)
- *Z*(*N*_NaCl_, *N, p, T*) : Isothermal-isobaric configurational partition function

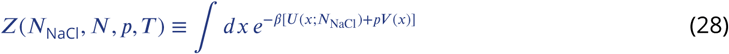
- Ξ(Δ*µ, N, p, T*) : Semigrand-isothermal-isobaric configurational partition function expressed as a sum over all *θ*

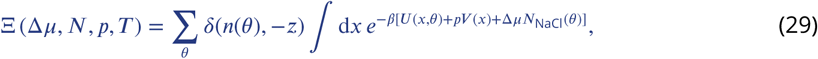

and expressed as a sum of number of ions and water molecules

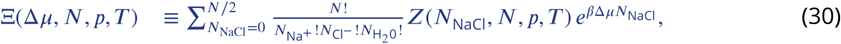

where 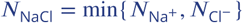 and 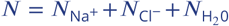. The upper bound of the summation— valid when *z* = 0 and *N* is even—is required as two water molecules are removed for every *N*_NaCl_.
- *π*(*x, θ*; *N, p, T*, *µ*) : Semigrand-isothermal-isobaric probability density with charge neutrality constraint

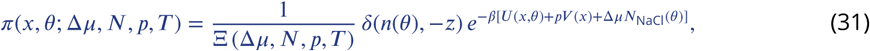

where the dependence of *π*(*x, θ*; Δ*µ, N, p, T*) on *z* is omitted for brevity
- ⟨*A*⟩_Δ*µ*_,_*N,p,T*_ : Expectation of *A*(*x, θ*) in (Δ*µ, N, p, T*) ensemble

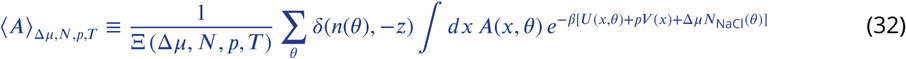
- 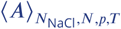: Expectation of *A*(*x*) in (*N*, *N, p, T*) ensemble

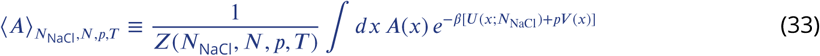

## Appendix 2

### Salt concentration in the thermodynamic limit

The purpose of this section is to derive an expression that relates the chemical potential to the salt concentration in a macroscopic saline reservoir (equation 19). This relationship is used in the calibration of our osmostat. The derivation will proceed by first, justifying the macroscopic concentration as the thermodynamic limit of the mean concentration, and second, rewriting the resultant expression in a manner that is amenable to computation.

#### The mean concentration in the thermodynamic limit

Following the definition of the concentration given in equation 20, the mean salt concentration in the semigrand ensemble considered here is given by

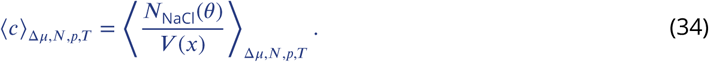

We seek an approximation to this expression that it is appropriate for large, macroscopic amounts of liquid saline. For brevity, all expectation values with respect to the thermodynamic ensemble (Δ*µ, N, p, T*) in this section will henceforth be abbreviated as ⟨·⟩.

The concentration is a function of two correlated random variables, the number of salt pairs *N*_NaCl_(*θ*) and the total volume *V* (*x*). A common way to approximate the expectation value, or mean, of a function of random variables is to perform a Taylor expansion about the mean of the arguments. The Taylor expansion (up to the second-order) of the function *g*(*a, b*) about the means ⟨*a*⟩ and ⟨*b*⟩, is

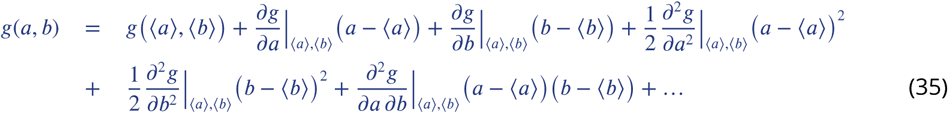

This expansion is particularly useful because the first order terms of the expanded mean ⟨*g*(*a, b*) ⟩ are zero i.e. ⟨*a* − ⟨*a*⟩ ⟩ = 0 and ⟨*b* − ⟨*b*⟩ ⟩ = 0. Hence, truncating the expansion to the second order leaves us with the approximation

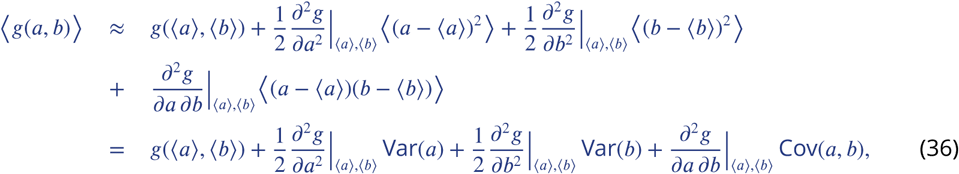

where Var(*a*) and Cov(*a, b*) denote the variance and covariance, respectively. Returning to the salt concentration, we relate *c* to the above with *g*(*N*_NaCl_, *V*) = *N*_NaCl_/*V*, and evaluate the partial derivatives to find that

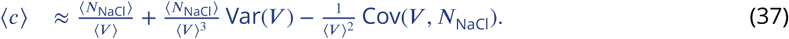

The leading term ⟨*N*_NaCl_⟩ /⟨*V*⟩ is the macroscopic expression that we seek. Thus, we require that the variance and covariance terms vanish in the thermodynamic limit. To show that they indeed do, we exploit the useful correspondence between partial derivatives and covariance in statistical thermodynamics. First, note that

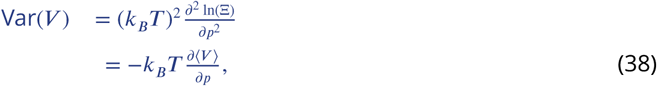

where Ξ ≡ Ξ(Δ*µ, N, p, T*) and is defined in equation 7. Also, note that

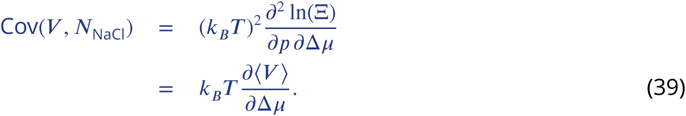

Second, we make use of the isothermal compressibility

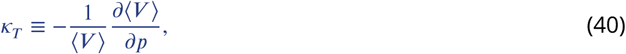

and introduce the isothermal susceptibility of the volume with respect to the chemical potential

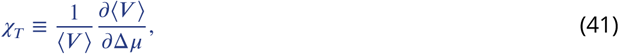

The susceptibilities *κ*_*T*_ and *χ*_*T*_ are bulk properties that measure the relative amount the volume of a system responds to changes in pressure and chemical potential, respectively. They are intensive quantities, such that they do not scale with the size of the system. These allow us to re-write the approximation of the mean concentration (equation 37) as

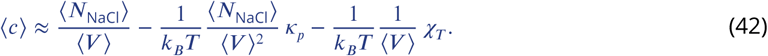

To proceed, note that in the second term, both *N*_NaCl_ and ⟨*V*⟩ are extensive, and rise in proportion to the total number of molecules in the system *N*. Thus, approximating the mean concentration as ⟨*N*_NaCl_⟩/ ⟨*V*⟩ incurs an error that is 𝒪(⟨*V*⟩^−1^), which tends to zero in the thermodynamic limit. We therefore define the macroscopic concentration of a saline reservoir as

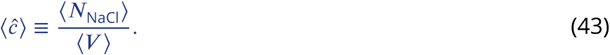

#### We require the macroscopic concentration to be amenable to computational analysis

While the expression for the macroscopic concentration above does not appear immediately useful, we now show how ⟨*ĉ*⟩ can be calculated for wide range of applied chemical potentials by pre-calculating the free energies to insert salt into a system, *f*(*N*_NaCl_) (≡ *f*(*N*_NaCl_, Δ*µ, N, p, T*)), and the average volume as a function of the number of salt pairs,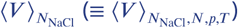.

To begin, it is useful to expand the definition of ⟨*N*_NaCl_⟩ given by equation 17 into

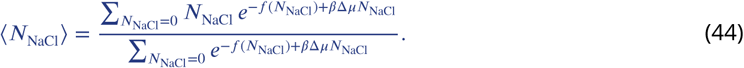

Next, we derive an expression for ⟨*V*⟩ that will cancel with the denominator of equation 44 when evaluating ⟨*ĉ*⟩. Using the representation of the semigrand density given by equation 8, the mean volume is given by

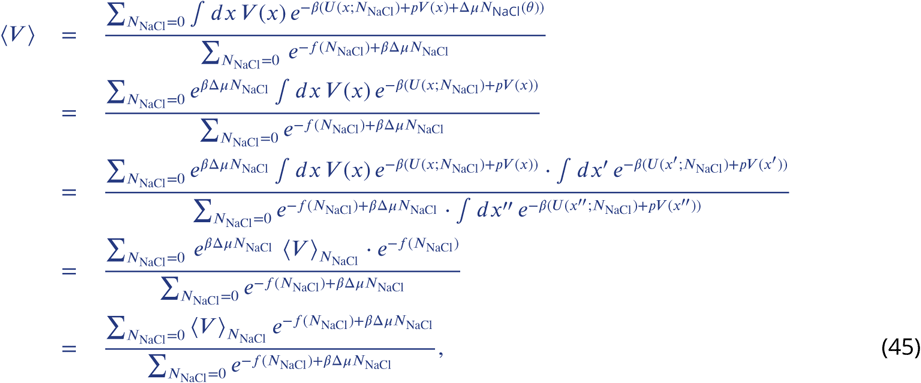

where the third and fourth line exploit the definition of the ensemble average for a fixed *N*_NaCl_. Inserting the expressions for the average number of salt pairs (equation 44) and the average volume (equation 45) into the macroscopic concentration (equation 43), we arrive at

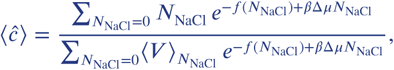

which is the same as equation 19 from the main text. Pertinently, the denominators in equations 44 and 45 have canceled, which greatly simplifies the evaluation of the macroscopic concentration for a given Δ*µ*.

### The magnitude of salt fluctuations

The concentration of salt fluctuates in osmostat simulations. This section briefly outlines how one would expect the magnitude of salt fluctuations to vary with the size of the system based on statistical mechanical principles. By differentiating equation 17, one can show that the variance of the number of salt pairs ⟨*N*_NaCl_⟩ is proportional to the gradient of *N*_NaCl_ with respect to the chemical potential Δ*µ*, specifically

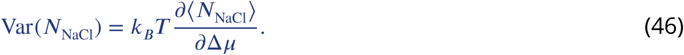

By dividing both sides by ⟨*N*_NaCl_⟩, i.e.

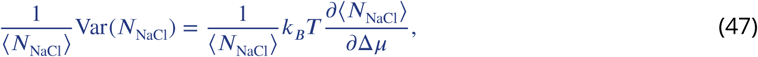

reveals that 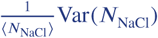 is proportional to the *relative* change in the mean of *N*_NaCl_ in response to altering the chemical potential. As the right-hand-side of the above equation is an intensive quantity, 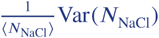 is also an intensive, implying that

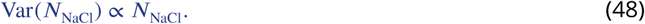

Therefore, the scale of the fluctuations in salt amount, as measured by the standard deviation, grows as ⟨*N*_NaCl_⟩^1/2^.

In contrast to the amount of salt, the size of the fluctuations of salt concentration *decreases* with the size of aqueous systems. Water is a highly incompressible fluid, such that small changes in pressure have a very small effect on the volume of aqueous systems. From equations 38 and 40, a low isothermal compressibility implies that the variance of the volume is small with respect to the mean volume (i.e. the relative variance). Assuming that the relative variance of the volume is smaller than the relative variance of the number of salt pairs, one can use the same approach as that of equation 35 to show that

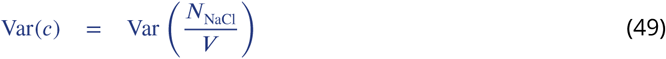

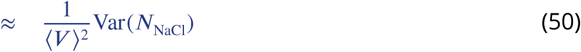

Using the fact that, for bulk-like water, 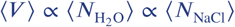 along with equation 48, we arrive at Var(*c*) ∼ ⟨*N*_NaCl_⟩^−1^ for systems with large amounts of water. Thus, the standard deviation of the salt concentration scales like 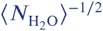 or ⟨*N*_NaCl_⟩^−1/2^ for a fixed chemical potential.

## Appendix 3

### Algorithmic implementation of the osmostat

This section describes the Metropolis-Hastings procedure from Saltswap [0.52] used to insert and delete salt. Insertion and deletion moves were enhanced with NCMC^32^. To describe its implementation of NCMC within SaltSwap, a more compressed notation is used compared to the original publication. For a more general and detailed exposition on NCMC, we refer readers to the original manuscript.

The osmostat move begins with the random choice of whether to insert or delete salt. The protocol is denoted Λ ∈ {Λ_insert_, Λ_delete_}, and the time reversed protocol is denoted 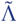, where 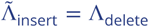 and 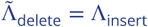. The probability to insert or delete a salt pair, *P* (Λ|*N*_NaCl_), depends on the number of salt molecules, *N*_NaCl_, in the system in the following way:

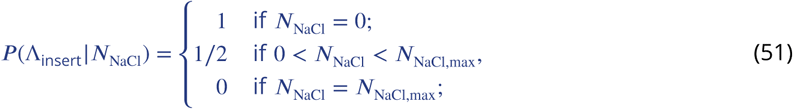

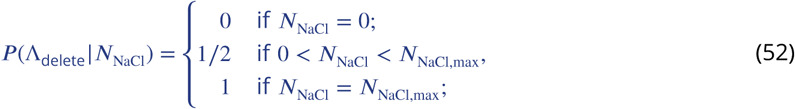

where for all simulations except the SAMS calibration simulations, 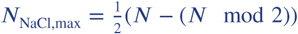 was chosen as two water molecules are required for the insertion of a Na^+^ and Cl^−^ pair. In the SAMS calibration simulations, *N*_NaCl,max_ was set to twenty. The particular choices of *P* (Λ_delete_|*N*_NaCl_) and *P* (Λ_insert_ |*N*_NaCl_) ensure that insertions are always attempted when there is no salt in the system, and deletions are always attempted when the number of salt pairs has reached maximum capacity.

For the insertion of salt, any two water molecules could be selected for transformation into Na^+^ and Cl^−^. Similarly, for the removal of salt, any Na^+^ ion and Cl^−^ ion could be selected for transformation into two water molecules. Formally, let *s*(*N*) denote the set {1, 2, …, *N*}, i.e. the set of indices for all water molecules and ions. For salt insertion, the index of candidate Na^+^ ion was a random uniform sample from the set {*i* ∈ *s*(*N*) : *θ*_*i*_ = 0} and the index of the Cl^−^ ion was a random uniform sample from the set {*j* ∈ *s*(*N*) : *θ*_*j*_ = 0, *i* ≠ *j*}. For salt removal, indices were selected randomly and uniformally from the sets {*i* ∈ *s*(*N*) : *θ*_*i*_ = +1} and {*j* ∈ *s*(*N*) : *θ*_*j*_ = −1}. As indices were chosen with equal probability within each set of possible candidates, the ratio of selection probabilities for molecule indices for forward and reverse protocols are given by

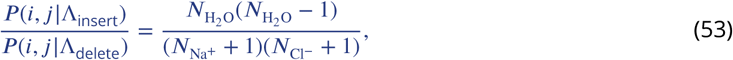

and

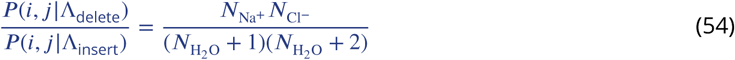

Following the choice of protocol and pair of molecules that would be transmuted, NCMC was used to enhance the efficiency of the insertion or deletion attempt. This implementation of NCMC consists of a fixed series of *perturbation* and *propagation* kernels over a fixed alchemical path. For both insertion and deletion moves, the alchemical path is a linear interpolation the nonbonded parameters of the water model and the ions. This particular alchemical path ensured that charge neutrality was maintained throughout the NCMC procedure.

The alchemical path is broken up into *T* segments that are uniformally spaced with respect to the nonbonded parameters. At state *t*, the configuration of the system will be denoted as *x*_*t*_ and the values of the nonbonded parameters for molecules *i* and *j* will be denoted as 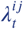. A single NCMC *step* corresponds to the application of the perturbation kernel followed by a the propagation kernel. When in state *t*, the perturbation kernel updates the nonbonded parameters 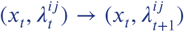, and the propagation kernel updates the configuration 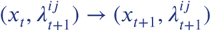. Each propagation kernel consists of *K* steps of Langevin dynamics using the parameters described in Simulation Details. A propagation kernel is also applied to the system before the first perturbation kernel to ensure the time symmetry of the protocol. The instantaneous change in the potential energy that results from the application of the perturbation kernel is recorded for each NCMC step and summed to produce the total work performed on the system by the protocol:

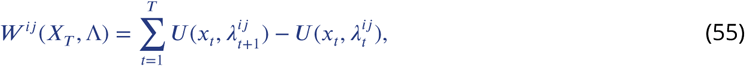

where the nonequilibrium trajectory *X*_*T*_ ≡ (*x*_0_, *x*_1_, …, *x*_*T*_). The difference between the protocol work and applied chemical potential Δ*µ*, along with the move proposal probabilities, determines whether a move is accepted or rejected. For the insertion of salt 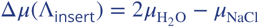, and for the deletion of salt 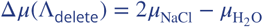. Attempts are accepted with the following probability

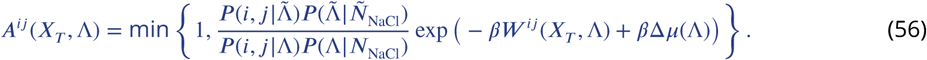

To preserve pathwise detailed balance, velocities were reversed upon acceptance. If a move is accepted, *θ*_*i*_ and *θ*_*j*_ are updated to reflect the new molecule identities.

### Pseudo-code for the NCMC osmostat with molecular dynamics

This section contains the pseudo-code of the production osmostat simulations.

#### Begin algorithm

Choose a macroscopic salt concentration *ĉ*.

Infer the chemical potential Δ*µ* by inverting equation 19.

Initialize position and velocity (*x*_0_, *v*_0_), state vector *θ*_0_, and maximum number of iterations *M*.

**for** *i* ∈ {1, 2, …, *M*} **do Sample conformations**

Perform 4 ps of Langevin integration with a fixed amount of salt:

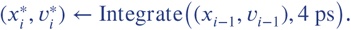

**Sample salt concentration**

Randomly select whether to add or remove salt as well as which molecules will be transmuted.

Define the trial state vector as *θ*^*^.

Define initial and final nonbonded parameters: (*q*_initial_, *σ*_initial_, *ϵ*_initial_) and (*q*_final_, *σ*_final_, *ϵ*_final_).

**procedure 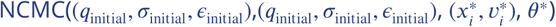**

Initialize variables, including protocol work *W* :

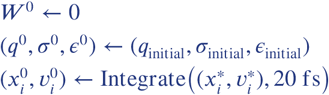

**for** *k* ∈ {1, 2, …, 1000} **do**

Linear interpolation of the nonbonded parameters:

*f* ^*k*^ = *k*/1000

**for all** atoms in the molecule **do**

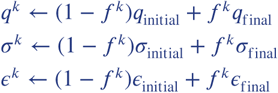

**end for**

Update the protocol work:

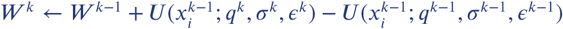

Propagate the system:

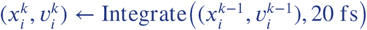

**end for**

Accept or reject using acceptance criterion *A*(*W* ^*k*^, Δ*µ, θ*^*^)

**if** Accept move **then**

Keep final positions and state vector but reverse velocities:

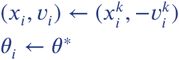

**else**

Return positions, velocities and the state vector to after equilibrium sampling:

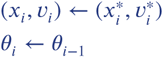

**end if**

**end procedure**

**end for**

**End algorithm**

## Appendix 4

### Validation: Ideal Mixing with the osmostat

In the Results section, Figure 4 *top left* indicates that the chemical potential has been properly calibrated, and Figure 6 shows that the osmostat produces samples that are concordant with physical-chemical intuition. In this section, we apply our osmostat to sample ideal mixing to provide further validation of the SaltSwap code base. Ideal mixing can be simulated with our osmostat by ensuring that salt insertion and deletion accrue no protocol work. This is implemented by using the same forcefield parameters for Na^+^ and Cl^−^ as the water model. As our osmostat also gives the ions the same mass as water, the “ions” sampled over in this section are identical to water except for their labeling.

To validate the sampling of the osmostat, we require an analytical relationship between the chemical potential Δ*µ* and the numbers of salt *N*_NaCl_ and water molecules 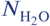. The chemical potential used in our osmostat is the difference between the chemical potential of water multiplied by two and Na^+^ and Cl^−^:

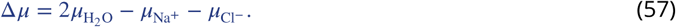

In order to relate Δ*µ* to *N*_NaCl_ and 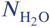, we will first consider a solution of water and ions in the (*N, p, T*) ensemble with fixed particle identities, and then relate the result to the (Δ*µ, N, p, T*) ensemble. For this fixed identity solution, let 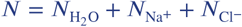 and 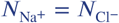. In the (*N, p, T*) ensemble, the chemical potential for a species *s* can be expressed as

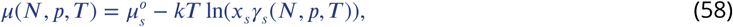

where 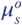 is the chemical potential of *s* in some reference state, *x*_*s*_ is the mole fraction of *s*, and *γ*_*s*_(*N, p, T*) is the activity coefficient of *s*. In general, the chemical potential is also dependent on the composition of the system. When Na^+^ and Cl^−^ have the same forcefield parameters and mass as water (i.e they are physically identical), the reference state and activity coefficients must be the same. So using equation 58 and 57 we have

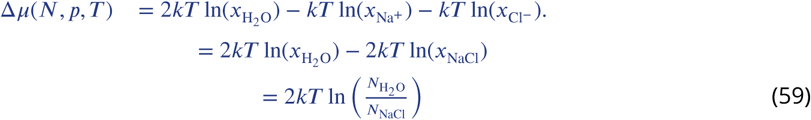

where the second line follows from the fact that there are equal numbers of Na^+^ and Cl^−^ ions. In the semigrand canonical (Δ*µ, N, p, T*) ensemble that is sampled by our osmostat, the chemical potential Δ*µ* is a controlled by the user. As this conjugate to the number of salt pairs, equation 59 will apply to the averages ⟨*N*_NaCl_⟩_Δ*µ*,N,p,T_ and 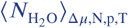, so that we have

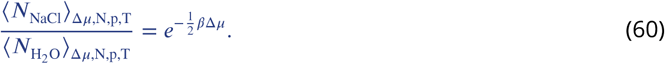

To test whether our osmostat correctly samples the average salt to water ratio given in equation 60, ideal mixing simulations were performed using SaltSwap on a small box of TIP3P water containing five hundred molecules for a range of chemical potentials. Ten thousand insertion and deletion attempts were made for salt pairs that had the same forcefield parameters as water. Only one perturbation step was used for the ideal NCMC insertion and deletion and the configuration of the system was not propagated during attempts. Figure 1 shows that there is excellent agreement between the relationship predicted by equation 60 and the simulation data.

**Appendix 4 Figure 1.**
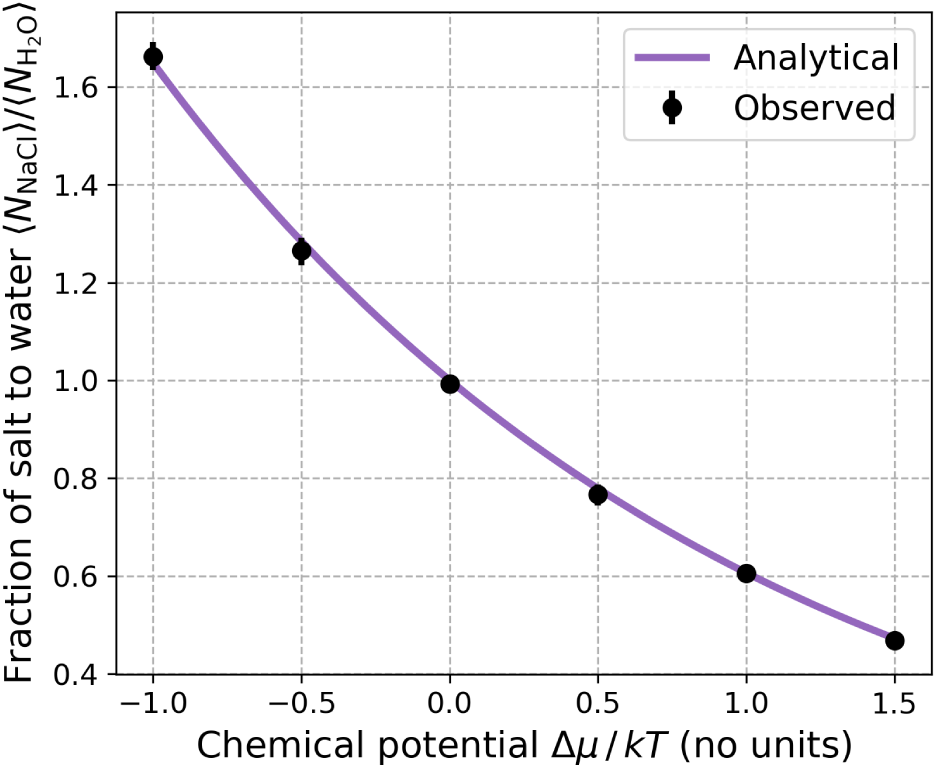
Validating the osmostat by comparing the observed average salt-water fractions to analytical values for ideal mixing. The relationship between the chemical potential and fraction of average number of salt pairs to water molecules is known exactly for ideal mixing, and is given by equation 60. Ideal mixing was implemented for the osmostat by giving the ions the same forcefield parameters as water. For each simulation at a chemical potential, the equilibration time and statistical inefficiency for the average number of salt pairs ⟨*N*_NaCl_⟩_Δ*µ*,N,p,T_ and water molecules 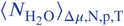 was determined using the timeseries module of pymbar^75^. The automatically determined equilibration times ranged from 361 and 723 insertion or deletion attempts. Effectively independent samples were extracted using the statistical inefficiency, and the means and 95% confidence intervals were estimated using bootstrap analysis.

It was also verified that the protocol work was effectively zero for the ideal NCMC transformations. While the protocol work should be exactly zero, the numerical imprecision of our implementation meant this could not always be achieved. The average protocol work for the transformations shown in Figure 1 (which were performed on a CPU Intel Core i7 with one perturbation step) was 1× 10^−7^ kT with a maximum absolute value of 8 × 10^−5^ kT. The NCMC protocol used throughout this study has one thousand perturbation steps and ten propagation steps per perturbation. With this protocol, the average protocol work was estimated using one thousand attempts on a GTX1080 GPU to be 2 × 10^−8^ kT with a maximum absolute value of 5 × 10^−4^ kT.

## Appendix 5

### Supplementary figures

**Appendix 5 Figure 1.**
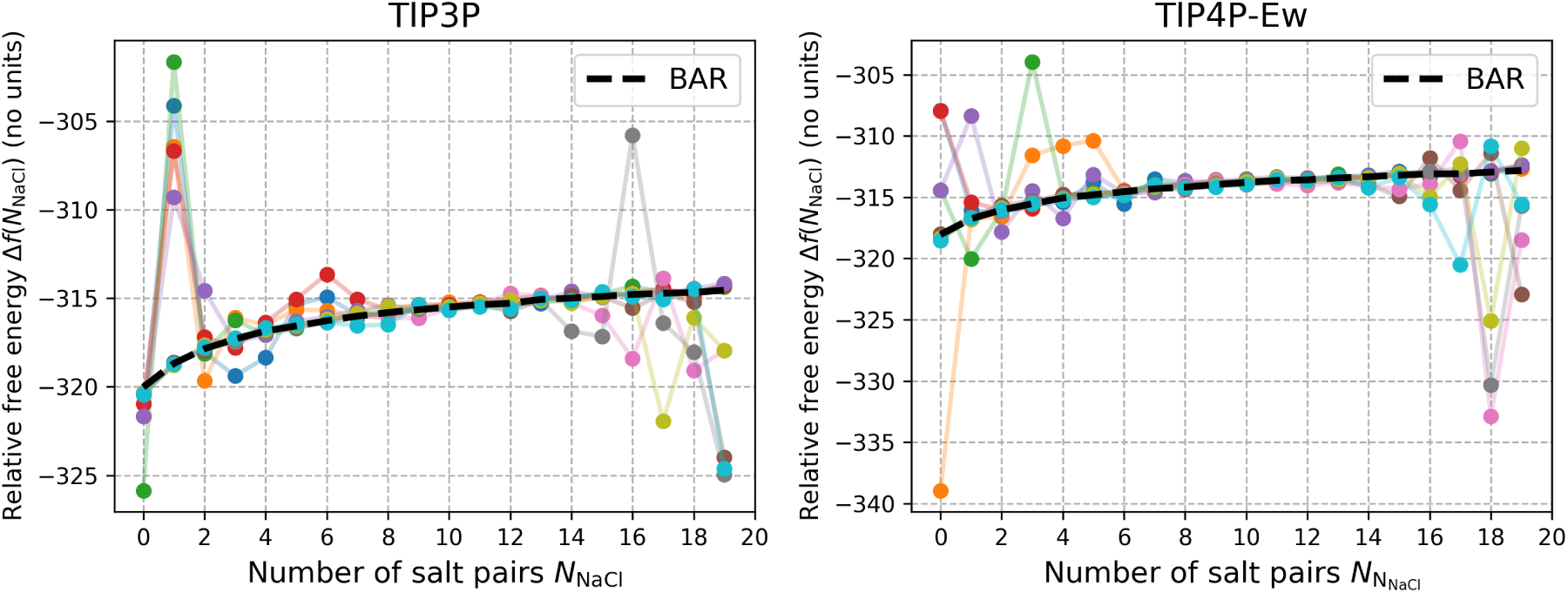
A Comparison of the salt insertion free energies as estimated by SAMS and BAR. The individual SAMS estimates from ten repeats of the relative free energy Δ*f*(*N*_NaCl_) to insert an Na^+^ and Cl^−^ and remove two water molecules in boxes of TIP3P (left) and TIP4P-Ew (right) for each SAMS simulations. Each color represents an estimate of Δ*f*(*N*_NaCl_) from each repeat. The relative free energy as calculated by BAR using all the SAMS simulation data is shown for reference (dotted black line). Five of the SAMS repeats were started with the maximum of 20 salt pairs in the system, and the other five started with none. The significant variation between the individual SAMS repeats is due to the rapid accumulation of the biasing potential in the initial stages of the algorithm. This biased the sampling away from the initial states of the simulations and prevented the uniform sampling over the salt numbers.

**Appendix 5 Figure 2.**
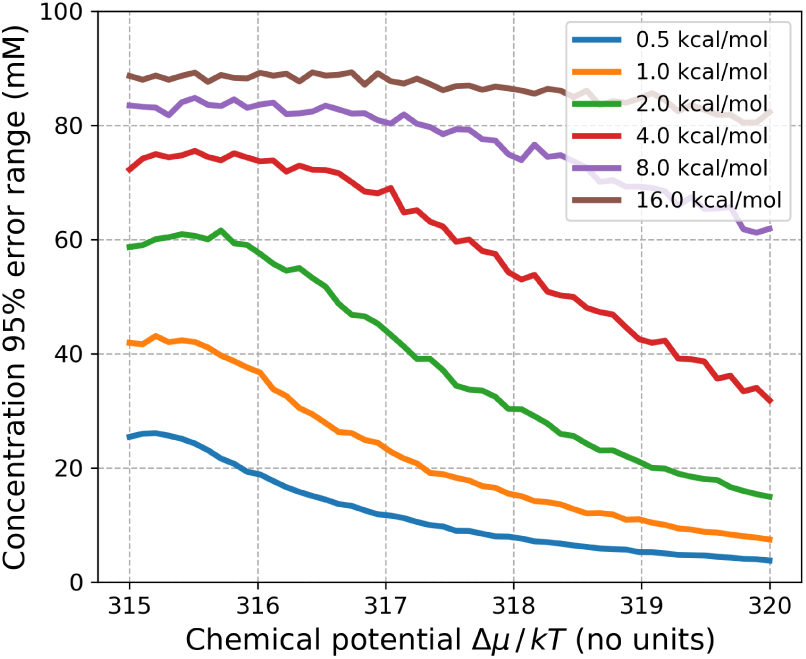
The statistical uncertainty of the predicted macroscopic concentration as a function of the chemical potential for different standard errors of the free energies *f*(*N*_NaCl_) in a box of 887 TIP3P water molecules. Using the data from the SAMS calibration simulations, Gaussian noise, with a mean of zero, was added to each estimated free energy *f*(*N*_NaCl_) *N* ∈ {0, 1, …, 20}, for a fixed values of 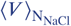. Three thousand noisy sample of *f*(*N*_NaCl_) *N* ∈ {0, 1, …, 20}, equation 19 were used to predict the macroscopic concentration for a range of chemical potentials. This figure shows the 95% confidence range of the resultant ensemble of concentrations for different standard deviations of the Gaussian noise about the free energies. One needs to evaluate the free energies *f*(*N*_NaCl_) to within 4 kcal/mol to achieve an error in the concentration that is no larger than roughly 80 mM. The tapering of the statistical error in the concentration at lower values of the chemical potential is due to maximum number of salt pairs used in the calibration (20), which limits that maximum concentration that can be predicted.

**Appendix 5 Figure 3.**
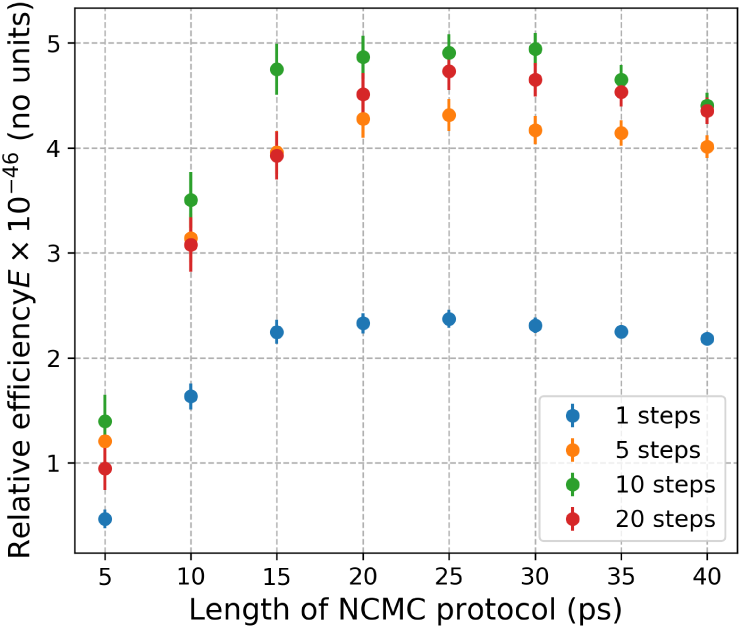
The relative efficiency of salt insertions/deletions in TIP3P water for different numbers of NCMC propagation steps between each perturbation step. Due to the manner in which the nonbonded parameters are updated in the SaltSwap code, it is faster—for a fixed protocol time-length—to perform multiple propagation steps for each perturbation (i.e. update of the nonbonded parameters) during an NCMC insertion/deletion attempt. More propagation steps limit the amount of communication between the CPU and GPU. However, for a fixed total protocol time-length, fewer perturbations increases the thermodynamic length each perturbation must traverse, which decreases the mean acceptance rate of the attempts. Thus, there is a (code-dependent) trade-off in the sampling efficiency between the number of perturbations and propagations steps. This figure shows the efficiency, defined by equation 25, for different numbers of propagation steps at different protocol time-lengths relative to the efficiency of instantaneous insertions and deletions. Ten propagation steps per perturbation step achieve the highest efficiencies, and so were used in all production osmostat simulations.

## References

[1] Moore, R. D.; Morrill, G. A. A possible mechanism for concentrating sodium and potassium in the cell nucleus. Biophys. J. 1976, 16, 527–533.

[2] Milo, R.; Phillips, R. Cell Biology by the Numbers, draft ed.; Garland Science, 2015; pp 127–130.

[3] Olaf S., A. Encyclopedia of Metalloproteins; Springer, 2013; pp 580–587.

[4] Le Rudulier, D.; Strom, A. R.; Dandekar, A. M.; Smith, L. T.; Valentine, R. C. Molecular biology of osmoregulation. Science 1984, 224, 1064–1068.

[5] Eil, R. et al. Ionic immune suppression within the tumour microenvironment limits T cell effector function. Nature 2016, 537, 539–543.

[6] Song, Y.; Gunner, M. R. Using multiconformation continuum electrostatics to compare chloride binding motifs in alpha-amylase, human serum albumin, and Omp32. J. Mol. biol. 2009, 387, 840–856.

[7] Southall, N. T.; Dill, K. A.; Haymet, A. D. J. A view of the hydrophobic effect. J. Phys. Chem. B 2002, 106, 521–533.

[8] Hribar, B.; Southall, N. T.; Vlachy, V.; Dill, K. A. How ions affect the structure of water. J. Am. Chem. Soc. 2002, 124, 12302–12311.

[9] Hofmeister, F. Zur lehre von der wirkung der salze: zweite mittheilung. Archiv Exp. Path. Pharm. 1888, 24, 247–260.

[10] Melander, W.; Horváth, C. Salt effects on hydrophobic interactions in precipitation and chromatography of proteins: An interpretation of the lyotropic series. Arch. Biochem. Biophys. 1977, 183, 200–215.

[11] Pegram, L. M.; Record, M. T. Hofmeister salt effects on surface tension arise from partitioning of anions and cations between bulk water and the air-water interface. J. Phys. Chem. B 2007, 111, 5411–5417.

[12] Hammarsten, O.; Chu, G. DNA-dependent protein kinase: DNA binding and activation in the absence of Ku. Proc. Natl. Acad. Sci. 1998, 95, 525–530.

[13] Lundbäck, T.; Härd, T. Salt dependence of the free energy, enthalpy, and entropy of nonsequence specific DNA binding. J. Phys. Chem. 1996, 100, 17690–17695.

[14] Misra, V. K.; Sharp, K. A.; Friedman, R. A.; Honig, B. Salt effects on ligand-DNA binding. J. Mol. Biol. 1994, 238, 245–263.

[15] Papaneophytou, C. P.; Grigoroudis, A. I.; McInnes, C.; Kontopidis, G. Quantification of the effects of ionic strength, viscosity, and hydrophobicity on protein-ligand binding affinity. ACS Med. Chem. Lett. 2014, 5, 931–936.

[16] Ong, W.; Kaifer, A. E. Salt effects on the apparent stability of the cucurbit [7] uril- methyl viologen inclusion complex. J. Org. Chem. 2004, 69, 1383–1385.

[17] Abraham, M. J.; Murtola, T.; Schulz, R.; Páll, S.; Smith, J. C.; Hess, B.; Lindahl, E. GROMACS: high performance molecular simulations through multi-level parallelism from laptops to supercomputers. SoftwareX 2015, 1-2, 19–25.

[18] Jo, S.; Kim, T.; Iyer, V. G.; Im, W. CHARMM-GUI: A web-based graphical user interface for CHARMM. J. Comput. Chem. 2008, 29, 1859–1865.

[19] Eastman, P.; Friedrichs, M. S.; Chodera, J. D.; Radmer, R. J.; Bruns, C. M.; Ku, J. P.; Beauchamp, K. A.; Lane, T. J.; Wang, L.-P.; Shukla, D.; et al., OpenMM 4: a reusable, extensible, hardware independent library for high performance molecular simulation. J. Chem. Theory Comput. 2013, 9, 461–469.

[20] Eastman, P.; Swails, J.; Chodera, J. D.; McGibbon, R. T.; Zhao, Y.; Beauchamp, K. A.; Wang, L.-P.; Simmonett, A. C.; Harrigan, M. P.; Stern, C. D.; et al., OpenMM 7: rapid development of high performance algorithms for molecular dynamics. PLOS Comput. Biol. 2017, 13, e1005659.

[21] Darden, T.; York, D.; Pedersen, L. Particle mesh Ewald: An *N* log *N* method for Ewald sums in large systems. J. Chem. Phys. 1993, 98, 10089–10092.

[22] Hub, J. S.; de Groot, B. L.; Grubmüller, H.; Groenhof, G. Quantifying artifacts in Ewald simulations of inhomogeneous systems with a net charge. J. Chem. Theory Comput. 2014, 10, 381–390.

[23] Wang, L.; Wu, Y.; Deng, Y.; Kim, B.; Pierce, L.; Krilov, G.; Lupyan, D.; Robinson, S.; Dahlgren, M. K.; Greenwood, J.; et al., Accurate and reliable prediction of relative ligand binding potency in prospective drug discovery by way of a modern free-energy calculation protocol and force field. J. Am. Chem. Soc. 2015, 137, 2695–2703.

[24] Ravishanker, G.; Auffinger, P.; Langley, D. R.; Jayaram, B.; Young, M. A.; Beveridge, D. L. In Reviews in Computational Chemistry; Lipkowitz, K. B., Boyd, D. B., Eds.; John Wiley & Sons, Inc.: Hoboken, NJ, USA, 1997; Vol. 11; pp 317–372.

[25] Ponomarev, S. Y.; Thayer, K. M.; Beveridge, D. L. Ion motions in molecular dynamics simulations on DNA. Proc. Natl. Acad. Sci. 2004, 101, 14771–14775.

[26] Rueda, M.; Cubero, E.; Laughton, C. A.; Orozco, M. Exploring the counterion atmosphere around DNA: what can be learned from molecular dynamics simulations? Biophys. J. 2004, 87, 800–811.

[27] Thomas, D. G.; Baker, N. A. GIBS: A grand-canonical Monte Carlo simulation program for simulating ion-biomolecule interactions. arXiv:Quantitative Biology 1704.05534.

[28] Vitalis, A.; Baker, N. A.; McCammon, J. A. ISIM: a program for grand canonical Monte Carlo simulations of the ionic environment of biomolecules. Mol. Simul. 2004, 30, 45–61.

[29] Shelley, J. C.; Patey, G. N. A configuration bias Monte Carlo method for ionic solutions. J. Chem. Phys. 1994, 100, 8265–8270.

[30] Lísal, M.; Smith, W. R.; Kolafa, J. Molecular simulations of aqueous electrolyte solubility: 1. The expanded-ensemble osmotic molecular dynamics method for the solution phase. J. Phys. Chem. B 2005, 109, 12956–12965.

[31] Moučka, F.; Lísal, M.; Skvor, J.; Jirsák, J.; Nezbeda, I.; Smith, W. R. Molecular simulation of aqueous electrolyte solubility. Osmotic ensemble Monte Carlo methodology for free energy and solubility calculations and application to NaCl. J. Phys. Chem. B 2011, 115, 7849–7861.

[32] Nilmeier, J. P.; Crooks, G. E.; Minh, D. D. L.; Chodera, J. D. Nonequilibrium candidate Monte Carlo is an efficient tool for equilibrium simulation. Proc. Natl. Acad. Sci. 2011, 108, E1009–E1018.

[33] Chen, Y.; Roux, B. Constant-pH hybrid nonequilibrium molecular dynamics–Monte Carlo simulation method. J. Chem. Theory Comput. 2015, 11, 3919–3931.

[34] Radak, B. K.; Chipot, C.; Suh, D.; Jo, S.; Jiang, W.; Phillips, J. C.; Schulten, K.; Roux, B. Constant-pH molecular dynamics simulations for large biomolecular Systems. J. Chem. Theory Comput. 2017,

[35] Tan, Z. Optimally adjusted mixture sampling and locally weighted histogram analysis. J. Comput. Graph. Stat. 2017, 26, 54–65.

[36] Lyubartsev, A. P.; Martsinovski, A. A.; Shevkunov, S. V.; Vorontsov-Velyaminov, P. N. New approach to Monte Carlo calculation of the free energy: method of expanded ensembles. J. Chem. Phys. 1992, 96, 1776–1783.

[37] Bennett, C. H. Efficient estimation of free energy differences from Monte Carlo data. J. Comput. Phys. 1976, 22, 245–268.

[38] Shirts, M. R.; Bair, E.; Hooker, G.; Pande, V. S. Equilibrium free energies from nonequilibrium measurements using maximum-likelihood methods. Phys. Rev. Lett. 2003, 91.

[39] Crooks, G. Excursions in statistical dynamics. Ph.D. thesis, 1999.

[40] Faller, R.; de Pablo, J. J. Constant pressure hybrid molecular dynamics–Monte Carlo simulations. J. Chem. Phys. 2002, 116, 55.

[41] Liu, J. Monte Carlo strategies in scientific computing; Springer Series in Statistics; Springer, 2008.

[42] Jorgensen, W. L.; Chandrasekhar, J.; Madura, J. D.; Impey, R. W.; Klein, M. L. Comparison of simple potential functions for simulating liquid water. J. Chem. Phys. 1983, 79, 926–935.

[43] Horn, H. W.; Swope, W. C.; Pitera, J. W.; Madura, J. D.; Dick, T. J.; Hura, G. L.; Head-Gordon, T. Development of an improved four-site water model for biomolecular simulations: TIP4P-Ew. J. Chem. Phys. 2004, 120, 9665–9678.

[44] Green, P. J. Reversible jump Markov chain Monte Carlo computation and Bayesian model determination. Biometrika 1995, 82, 711–732.

[45] Chodera, J. D.; Shirts, M. R. Replica exchange and expanded ensemble simulations as Gibbs sampling: simple improvements for enhanced mixing. J. Chem. Phys. 2011, 135, 194110.

[46] Speidel, J. A.; Banfelder, J. R.; Mezei, M. Automatic control of solvent density in grand canonical ensemble Monte Carlo simulations. J. Chem. Theory Comput. 2006, 2, 1429–1434.

[47] Benavides, A. L.; Aragones, J. L.; Vega, C. Consensus on the solubility of NaCl in water from computer simulations using the chemical potential route. J. Chem. Phys. 2016, 144, 124504.

[48] Ross, G. A.; Bodnarchuk, M. S.; Essex, J. W. Water sites, networks, and free energies with grand canonical Monte Carlo. J. Am. Chem. Soc. 2015, 137, 14930–14943.

[49] Ross, G. A.; Bruce Macdonald, H. E.; Cave-Ayland, C.; Cabedo Martinez, A. I.; Essex, J. W. Replica exchange and standard state binding free energies with grand canonical Monte Carlo. J. Chem. Theory Comput. 2017,

[50] Marinari, E.; Parisi, G.; Roma, S.; Vergata, T. Simulated tempering: a new Monte Carlo scheme. 1992,

[51] Wang, F.; Landau, D. P. Efficient, multiple-range random walk algorithm to calculate the density of states. Phys. Rev. Lett. 2001, 86, 2050–2053.

[52] Liang, F. A generalized Wang-Landau algorithm for Monte Carlo computation. J. Am. Stat. Assoc. 2005, 100, 1311–1327.

[53] Fukuda, I.; Nakamura, H. Non-Ewald methods: theory and applications to molecular systems. Biophys. Rev. 2012, 4, 161–170.

[54] Lieb, E. H.; Lebowitz, J. L. The constitution of matter: existence of thermodynamics for systems composed of electrons and nuclei. Adv. Math. 1972, 9, 316–398.

[55] Neal, R. M. Annealed Importance Sampling. arXiv:physics 1998, 9803008.

[56] Karagiannis, G.; Andrieu, C. Annealed importance sampling reversible jump MCMC algorithms. J. Comput. Graph. Stat. 2013, 22, 623–648.

[57] Sivak, D. A.; Chodera, J. D.; Crooks, G. E. Using nonequilibrium fluctuation theorems to understand and correct errors in equilibrium and nonequilibrium simulations of discrete Langevin dynamics. Phys. Rev. X 2013, 3, 011007.

[58] Wagoner, J. A.; Pande, V. S. Reducing the effect of Metropolization on mixing times in molecular dynamics simulations. J. Chem. Phys. 2012, 137, 214105.

[59] Sohl-Dickstein, J. Hamiltonian Monte Carlo with reduced momentum flips. arXiv:physics 2012, 1205.1939.

[60] Chen, Y.; Roux, B. Efficient hybrid non-equilibrium molecular dynamics - Monte Carlo simulations with symmetric momentum reversal. J. Chem. Phys. 2014, 141, 114107.

[61] Elena, A.; Bou-Rabee, N.; Reich, S. A comparison of generalized hybrid Monte Carlo methods with and without momentum flip. J. Comput. Phys. 2009, 228, 2256–2265.

[62] Leimkuhler, B.; Matthews, C. Molecular Dynamics: With Deterministic and Stochastic Numerical Methods; Springer, 2015.

[63] OpenMMTools 0.11.1. https://github.com/choderalab/openmmtools/releases/tag/0.11.1, 07-06-2017.

[64] Amber 14 benchmark archive. http://ambermd.org/Amber14_Benchmark_Suite.tar.bz2, 07-06-2017.

[65] Maier, J. A.; Martinez, C.; Kasavajhala, K.; Wickstrom, L.; Hauser, K. E.; Simmerling, C. ff14SB: improving the accuracy of protein side chain and backbone parameters from ff99SB. J. Chem. Theory Comput. 2015, 11, 3696–3713.

[66] PDBFixer. https://github.com/pandegroup/pdbfixer, 07-06-2017.

[67] Zgarbová, M.; Šponer, J.; Otyepka, M.; Cheatham, T. E.; Galindo-Murillo, R.; Jurečka, P. Refinement of the sugar- phosphate backbone torsion beta for AMBER force fields improves the description of Z- and B-DNA. J. Chem. Theory Comput. 2015, 11, 5723–5736.

[68] Joung, I. S.; Cheatham, T. E. Determination of alkali and halide monovalent ion parameters for use in explicitly solvated biomolecular simulations. J. Phys. Chem. B 2008, 112, 9020–9041.

[69] Barth, E.; Kuczera, K.; Leimkuhler, B.; Skeel, R. D. Algorithms for constrained molecular dynamics. J. Comput. Chem. 1995, 16, 1192–1209.

[70] Eastman, P.; Pande, V. S. Constant constraint matrix approximation: a robust, parallelizable constraint method for molecular simulations. J. Chem. Theory Comput. 2010, 6, 434–437.

[71] Miyamoto, S.; Kollman, P. A. Settle: an analytical version of the SHAKE and RATTLE algorithm for rigid water models. J. Comput. Chem. 1992, 13, 952–962.

[72] Chodera, J. D. A simple method for automated equilibration detection in molecular simulations. J. Chem. Theory Comput. 2016, 12, 1799–1805.

[73] van der Walt, S.; Colbert, S. C.; Varoquaux, G. The NumPy array: a structure for efficient numerical computation. Comput. Sci. Eng. 2011, 13, 22–30.

[74] Jones, E.; Oliphant, T.; Peterson, P. SciPy 0.19.1. http://www.scipy.org, 06-23-2017.

[75] pymbar 3.0.1. https://github.com/choderalab/pymbar, 2-3-2017.

[76] McGibbon, R. T.; Beauchamp, K. A.; Harrigan, M. P.; Klein, C.; Swails, J. M.; Hernández, C. X.; Schwantes, C. R.; Wang, L.- P.; Lane, T. J.; Pande, V. S. MDTraj: a modern open library for the analysis of molecular dynamics trajectories. Biophys. J. 2015, 109, 1528–1532.

[77] Humphrey, W.; Dalke, A.; Schulten, K. VMD: visual molecular dynamics. J. Mol. Graph. 1996, 14, 33–38.

[78] Hunter, J. D. Matplotlib: A 2D graphics environment. Comput. Sci. Eng. 2007, 9, 90–95.

[79] Jorgensen, W. L.; Blake, J. F.; Buckner, Free energy of TIP4P water and the free energies of hydration of CH4 and Cl- from statistical perturbation theory. Chem. Phys. 1989, 129, 193–200.

[80] Marcus, Y. Thermodynamics of solvation of ions. Part 5. Gibbs free energy of hydration at 298.15 K. J. Chem. Soc., Faraday Trans. 1991, 87, 2995–2999.

[81] Noyes, R. M. Thermodynamics of Ion Hydration as a Measure of Effective Dielectric Properties of Water. J. Am. Chem. Soc. 1962, 84, 513–522.

[82] Schmid, R.; Miah, A. M.; Sapunov, V. N. A new table of the thermodynamic quantities of ionic hydration: values and some applications (enthalpy-entropy compensation and Born radii). Phys. Chem. Chem. Phys. 2000, 2, 97–102.

[83] Sastre de Vicente, M. E. The concept of ionic strength eighty years after its introduction in chemistry. J. Chem. Educ. 2004, 81, 750–753.

[84] Crooks, G. E. Measuring thermodynamic length. Phys. Rev. Lett. 2007, 99, 100602.

[85] Minh, D. D. L.; Chodera, J. D. Estimating equilibrium ensemble averages using multiple time slices from driven nonequilibrium processes: Theory and application to free energies, moments, and thermodynamic length in single- molecule pulling experiments. J. Chem. Phys. 2011, 134, 024111.

[86] Sivak, D. A.; Crooks, G. E. Thermodynamic metrics and optimal paths. Phys. Rev. Lett. 2012, 108, 190602.

[87] Rotskoff, G. M.; Crooks, G. E. Optimal control in nonequilibrium systems: dynamic Riemannian geometry of the Ising model. Phys. Rev. E 2015, 92, 060102.

[88] Gingrich, T. R.; Rotskoff, G. M.; Crooks, G. E.; Geissler, P. L. Near-optimal protocols in complex nonequilibrium transformations. Proc. Natl. Acad. Sci. 2016, 113, 10263–10268.

[89] Webb, B. A.; Chimenti, M.; Jacobson, M. P.; Barber, D. L. Dysregulated pH: a perfect storm for cancer progression. Nat. Rev. Cancer 2011, 11, 671–677.

[90] Spahn, V.; Del Vecchio, G.; Labuz, D.; Rodriguez-Gaztelumendi, A.; Massaly, N.; Temp, J.; Durmaz, V.; Sabri, P.; Reidel-bach, M.; Machelska, H.; et al., A nontoxic pain killer designed by modeling of pathological receptor conformations. Science 2017, 355, 966–969.

[91] Salvatier, J.; Wiecki, T.; Fonnesbeck, C. Probabilistic programming in Python using PyMC3. PeerJ Computer Science 2016, 2, e55.

[92] Andrieu, C.; Thoms, J. A tutorial on adaptive MCMC. Stat. Comput. 2008, 18, 343–373.

[93] Haario, H.; Saksman, E.; Tamminen, J. An adaptive Metropolis algorithm. Bernoulli 2001, 7, 223–242.

[94] Atkinson, C.; Mitchell, A. F. Rao’s distance measure. Sankhyā Series A 1981, 43, 345–365.

[95] Radak, B. K.; Roux, B. Efficiency in nonequilibrium molecular dynamics Monte Carlo simulations. J. Chem. Phys. 2016, 145, 134109.

